# Molecular aging is the main driver of Parkinson’s Disease

**DOI:** 10.1101/2025.08.27.672359

**Authors:** Verena Bopp, Jaehyun LeeBae, Patrick Oeckl, Julia K. Kühlwein, Veselin Grozdanov, Martin Kiechle, Benjamin Mayer, Bettina Möhrle, Hartmut Geiger, Karin M. Danzer

## Abstract

Aging as well as the presence of α-synuclein (α-syn) oligomers in the brain are indisputably linked to Parkinson’s disease (PD). A central concept of geroscience is that the biological processes of aging drive the onset of aging-associated diseases. The extent to which the biological processes of aging directly contribute to PD and the inter-relationship with α-syn oligomers for the onset of PD symptoms remains unclear. Using an inducible α-syn oligomer mouse model of PD, we demonstrate that the induction of PD associated α-syn oligomers for the same timespan caused PD associated symptoms only in aged, but not in young mice. Biochemical studies revealed that α-syn oligomer formation precedes motor decline in these aged mice, and age together with α-syn expression determine the motor phenotype. Single-nucleus RNA sequencing (snRNA-seq) identified a PD disease signature that was particularly linked to basal ganglia neurons (BGNs) and was in part shared with an aging transcriptional signature. PD symptoms, as well as the PD Signature, were significantly altered by a short-term pharmacological attenuation of the activity of the small RhoGTPase CDC42 in already aged animals with PD symptoms. Attenuation of activity of CDC42 is known to target the general biological processes of aging. Interestingly, the intervention did not affect the amount of α-syn oligomers in the animals, while still improving phenotypes. Together, the data demonstrates that the biological processes of aging are a major causative driver for the onset of PD in the α-syn model of PD.

## Introduction

Aging is the major risk factor for Parkinson’s disease (PD). The onset of symptoms typically occurs between the ages of 55 and 65 (Silkis 2001). The symptoms include motor disturbances but also non-motoric symptoms like sleep disturbances and cognitive impairment. The severity of symptoms tends to worsen with age. However, the age at disease onset can vary by decades and this variability in age of symptom onset significantly influences the progression of the disease (Raket et al. 2022).

One of the central molecular players implicated in the pathogenesis of PD is α-synuclein (α-syn), a presynaptic neuronal protein that is crucial for synaptic vesicle regulation. Under pathological conditions, α-syn misfolds and aggregates into oligomeric and fibrillar forms, and will by this means contribute to the neurodegenerative processes observed in PD (Ingelsson 2016). The accumulation of α-syn aggregates in mouse models for PD is increased in aged mice (Li et al. 2024; Kiechle et al. 2019) and there is more α-syn in the soma of neurons of the substantia nigra pars compacta in aged humans (Chu and Kordower 2007). A range of symptoms that are associated with PD, including motor and cognitive symptoms, may also arise due to aging and co-pathologies.

A central concept of geroscience is that the biological processes of aging drive the onset of aging-associated diseases. Whether molecular and cellular processes that underlay aging contribute causatively to PD initiation though is still not known (Martirosyan et al. 2024). We still cannot answer the fundamental question, whether the aged organism is more vulnerable to a disease triggering event, like α-syn aggregation, so that the disease develops primarily only upon aging, or whether for example α-syn oligomer deposits simply requires a distinct, long time to develop into disease phenotypes. The second scenario would mean that only the time span is important and thus independent of molecular or cellular changes that are associated with normal aging.

Aging at the cellular level refers to the progressive decline in the physiological function of cells, leading to reduced capacity for repair, replication, and response to stress. This decline is influenced by various interconnected biological processes and mechanisms. In addition to the progressive accumulation of molecular damage inside cells, aging is associated with an overall decrease in proteasome activity, impaired autophagy, mitochondrial dysfunction, rearrangements of the cytoskeleton and neuroinflammation, which are precisely the signaling pathways that are also dysregulated in neurodegenerative disease (Vanni et al. 2020). Additionally, aging primarily leads to impairments of intracellular clearance mechanisms, especially in the ventral substantia nigra which in turn increases vulnerability to neurodegeneration. An age-related association between the increase in α-syn and the decrease in nigral tyrosine hydroxylase has been demonstrated (Chu and Kordower 2007).

In recent years, advanced methodologies such as single-cell sequencing have facilitated a more detailed analysis of different cell types and states during disease progression and aging. This has opened new potential avenues for combating PD (Smajić et al. 2022), provided detailed resources for various cell types affected by PD or aging (Martirosyan et al. 2024; Ximerakis et al. 2019), and led to the discovery of new, disease- or age-relevant rare cell types (Kamath et al. 2022).

In this study, we utilized a well-characterized α-syn oligomer PD mouse model (Kiechle et al. 2019) to determine the age at which induction of α-syn oligomerization results in PD phenotypes. We further dissect α-syn oligomer and motor phenotypes in relation to time span and aging and study on a biochemical and single cell transcriptomics level which cellular pathways fuel PD disease development and/or aging.

## Results

For the onset of PD, aging is the major risk factor. We have previously described a transgenic α-syn PD mouse model in which the time-dependent accumulation of α-syn oligomers upon aging of the mice was accompanied by neuronal cell loss and diminished motoric abilities (Kiechle et al. 2019). The question whether the changes upon aging of the mice determined, in combination with α-syn oligomer expression, the motoric impairment or if simply a time-span of α-syn oligomer exposure resulted in the decline in motor function could not be determined. We therefore used an inducible PD mouse model based on a human α-syn protein complementation system expressing α-syn fused to halves of Gaussia luciferase (called S1/S2 model). The expression of this non-bioluminescent S1/S2 construct is neuron-specific and inducible. S1/S2 fragments are reconstituted when brought together by S1-S2 interactions (Tet Off System) (**Fig.1 A**).

**Figure 1:**
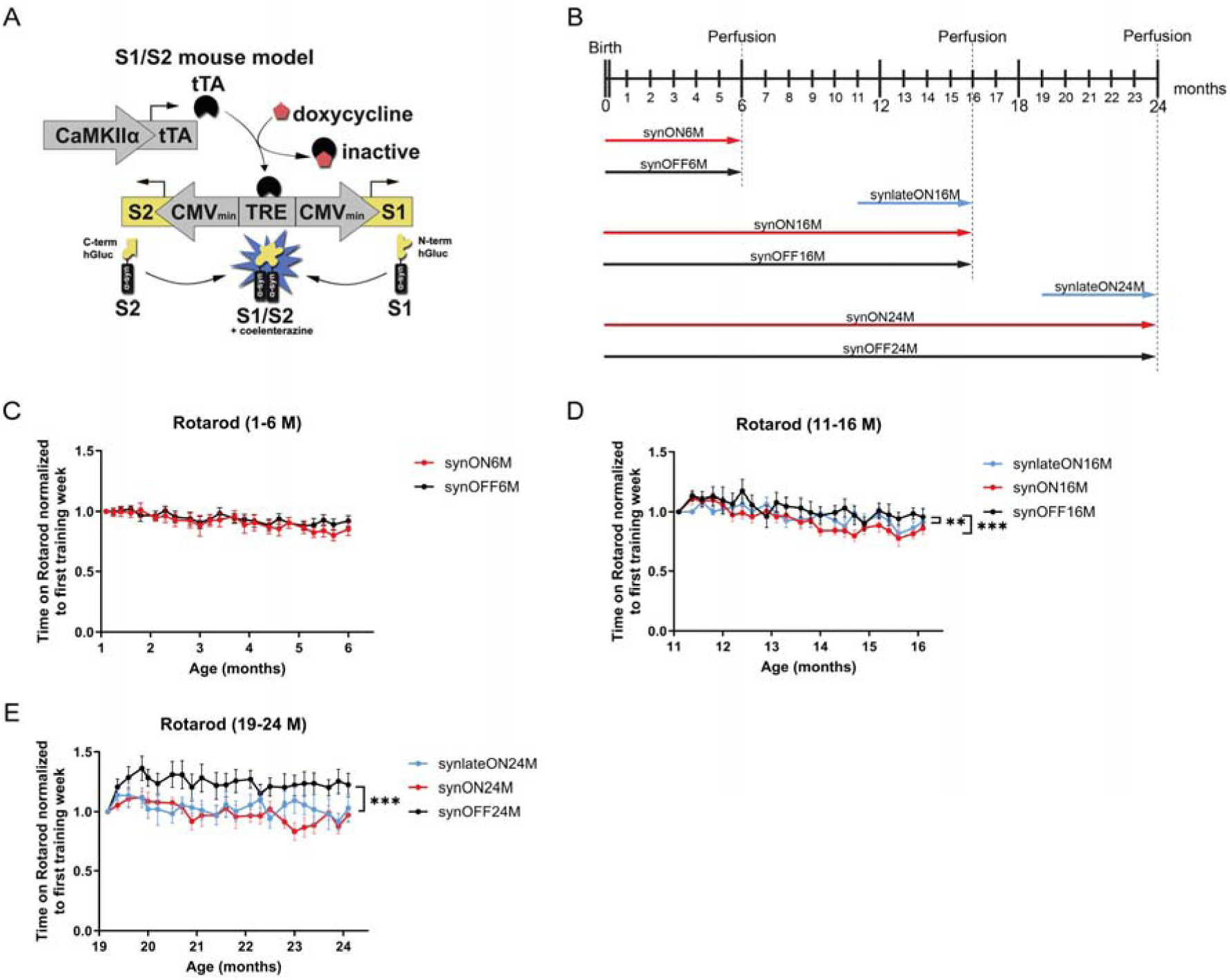
(A) Schematic principle of the S1/S2 PD mouse model which includes a protein complementation assay (PCA) expressing α-syn coupled to either the N- or C-terminal part of a Gaussia luciferase. The expression takes place under the neuron-specific CamKIIα promoter and is modeled as a Tet-off system driven by doxycycline. Figure modified from Kiechle et al., 2019. (B) Overview of all different S1/S2 mouse cohorts which differ in their age (6, 16 and 24 M) and length of S1/S2 expression (short-, long- and non-expressing). (C-E) Accelerating Rotarod was used to determine motor balance and coordination of all S1/S2 mouse cohorts, comparing short-, long- and non-expressing animals with (C) 1-6 M, (D) 11-16 M and (E) 19-24 M of age by measuring the latency to fall in seconds (s) (n=12 male and female mice, repeated-measures two-way ANOVA, ***p<0.001, **p<0.01; data: mean ± SEM).

To determine a potential vulnerable age for S1/S2 oligomer formation and/or motoric decline, we induced S1/S2 oligomer expression for only 5 months (M) at 11 and 19 M of age (synlateON16M and synlateON24M) and performed Rotarod testing. These groups were compared to mice with lifelong S1/S2 expression (synON) or no expression at all (synOFF) at the ages of 6, 16 and 24 M (**Fig.1 B**). Thus, our experimental S1/S2 mouse groups differ in their age and length of S1/S2 expression (**Fig.1 B**). Young synON and synOFF mice between 1 and 6 M of age showed no significant differences in their motoric abilities (**Fig.1 C**). In contrast, mice expressing S1/S2 for only 5 M starting at 11 M of age (synlateON16M) as well as mice expressing S1/S2 throughout their entire lives (synON16M) exhibited significantly reduced motor performance between 11 and 16 M of age compared to control animals (synOFF16M) (**Fig.1 D**). The progressive decline in motoric abilities became even more pronounced between 19 and 24 M of age in the S1/S2 expressing animals in which expression was only induced at 19 M of age (synlateON24M) throughout their entire lives (synON24M) compared to controls (**Fig.1 E**). Notably, at 24 M of age there was no difference in the motoric phenotype between the synlateON and synON condition. These results suggest that age, rather than the duration of oligomer exposure, determines the severity of the S1/S2- induced motoric phenotype with synlateON24M and synON24M animals showing the most significant motoric impairment compared to controls.

To determine whether the motoric alterations correlate with the total amount of expressed S1/S2, we performed capillary based Simple Western with anti-α-syn antibody. While the expression levels of S1/S2 between the groups slightly varied, none of the comparisons reached statistical significance (**Fig.2 A, B**). To gain insight into the various oligomer species, we performed Size-Exclusion Chromatography (SEC) with subsequent luciferase activity measurement in each fraction. Short expressing mice at the age of 16 M (synlateON16M) revealed a similar heterogeneous profile of S1/S2 oligomers as S1/S2 expressing mice from birth (synON16M) (**Fig.2 D**). In contrast 24 M old mice differed in their oligomer profile depending on whether S1/S2 oligomers were present since birth (synON24M) or only for 5 M (synlateON24M). This was also reflected in the total luciferase activity with a significantly higher area under the curve (AUC) for the life-long expressing animals at 24 M of age (**Fig.2 F**). A detailed analysis of the SEC profile separating the profile according to the two main peaks (void volume peak with sizes > 430 kDa and a peak reflecting 8-16-mers, **Fig.2 G-H**) revealed that 8-16-mers S1/S2 oligomers were enriched in synON24 animals compared to synlateON24 mice. Additionally, the void volume peak, containing aggregates larger than 430 kDa showed a tendency to increase for the synON24M animals compared to synlateON24M (**Fig.2 G-H**).

**Figure 2:**
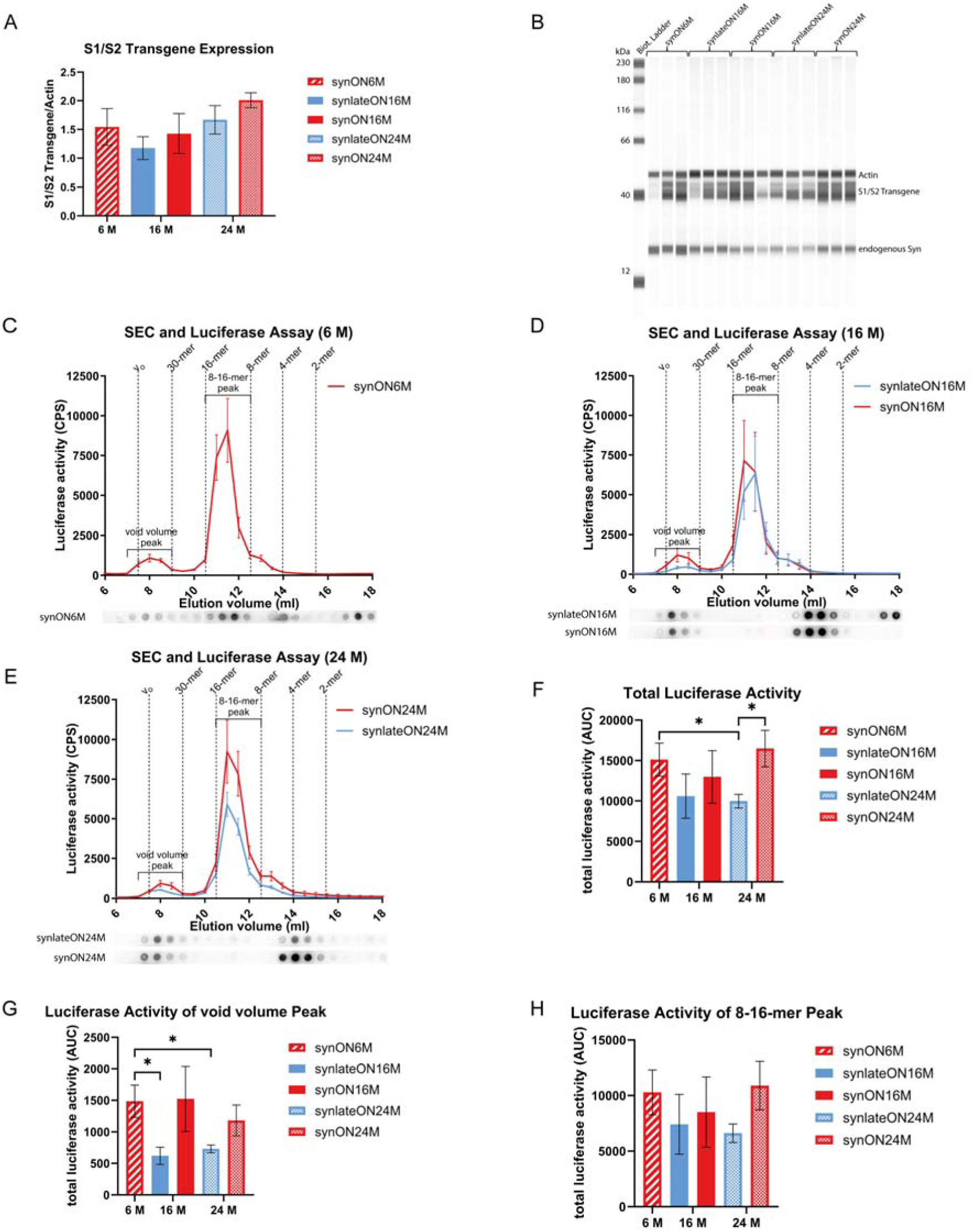
(A)-(B) Determination of S1/S2 transgene and endogenous α-syn expression of full-brain lysates from S1/S2 mice by capillary-based Western blot analysis (Simple Western) using the Jess system and normalization to Actin (anti-α-syn Syn1 antibody and anti-Actin antibody) (student’s t-test, ***p<0.001; data: mean ± SEM). (C)-(H) Gaussia luciferase activity and dot blot analysis (anti-α-syn 15G7 antibody) of SEC fractions of full-brain lysates from S1/S2 animals of (C) 6 M, (D) 16 M and (E) 24 M old mice. The graphs illustrate the mean ± SEM of n=5-6 animals per group. There is a clear enrichment of 8-16-mer species in all mice groups. (F)-(H) Quantification of the AUC of respective total luciferase activity of SEC fractions (one-way ANOVA with Tukey’s post hoc test, *p<0.05; data: mean ± SEM).

Since the motoric impairment in PD mice was absent at 6 M, we speculated that a lower level of S1/S2 oligomers would be present in this age group. However, contrary to our assumption, 6 M old S1/S2 expressing mice already exhibited a similar S1/S2 oligomer profile to that of 16 or 24 M old PD mice (**Fig.2 C**). Together, these results suggest that S1/S2 oligomer formation precedes motoric decline in PD mice, with the age of the PD mice mainly determining the motoric phenotype. In contrast, the duration and timing of S1/S2 expression mainly affect oligomer load.

The data demonstrated so far support that the biological processes of aging are a major causative driver for the onset of PD in the α-syn PD model. Aging has been associated with an elevated level of the activity of the small RhoGTPase Cell division control protein 42 homolog (CDC42) (L. Wang et al. 2007; Florian et al. 2012). This increase in CDC42 activity is causative for aging-associated phenotypes in multiple tissues and types of cells (Umbayev et al. 2023). Also in brain, there is significant increase in the activity of CDC42 upon aging (L. Wang et al. 2007). Moreover, a novel association of loss of function variants of a CDC42 activating gene *ITSN1* and PD have been found (Skuladottir et al. 2024). Attenuation of CDC42 activity via the pharmacological compound CASIN (CDC42 activity specific inhibitor) in already aged mice increases life span (Florian et al. 2020). Attenuation of the activity of CDC42 therefore targets basic cellular and molecular pathways of aging (Umbayev et al. 2023). We therefore tested whether CDC42 inhibition by CASIN might potentially rescue the PD associated motoric decline in α-syn expressing aged mice and administered it around the age of PD onset (12 M of age) and after age of onset (20 M of age). CASIN was administered in two rounds, each consisting of four consecutive daily doses with a one-week interval between rounds, for a total of eight applications. Indeed, systemic treatment with CASIN (similar to the treatment regimen that results in lifespan extension, **Fig.3 A**) ameliorated the motoric decline at the age 11-16 M, and very pronounced, at 21-24 M range of age (**Fig.3 B,C**). Interestingly, CASIN treatment did not affect the amount of α-syn oligomers. Aged mice treated with CASIN improved their motor function but had similar amounts of oligomers compared to aged, non-treated animals (**Fig.3 D-G**). These data confirm that while oligomers are necessary to develop PD symptoms, they are not sufficient, like the presence of oligomers in young animals (**Fig.2 C**). They further confirm that attenuating aging-related changes can reverse PD phenotypes in this α-syn model without affecting the α-syn oligomer load.

**Figure 3:**
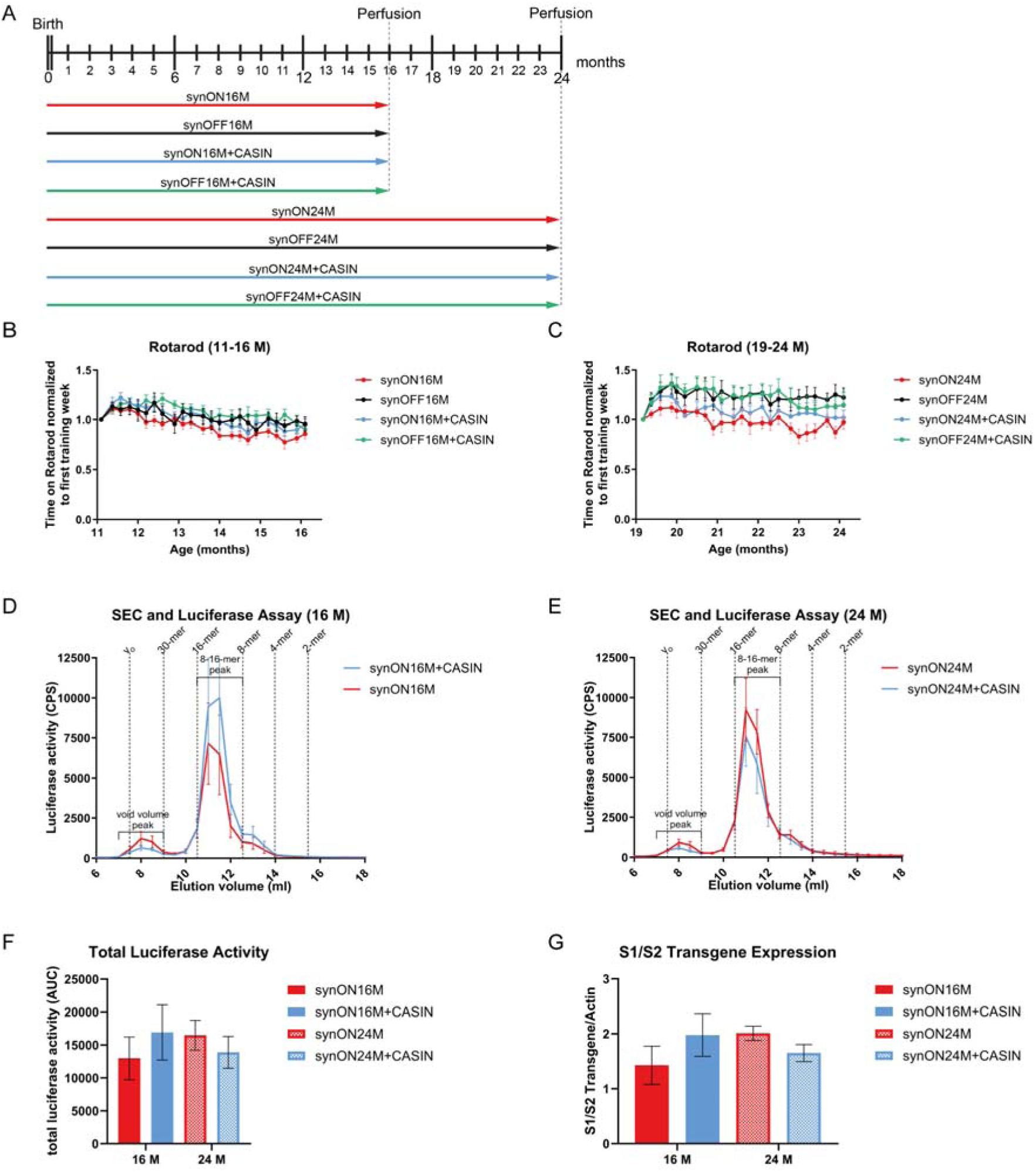
(A) Overview of the different S1/S2 mouse cohorts for CASIN treatment. They are categorized by age (16 and 24 M), levels of S1/S2 expression (ON vs OFF) and treatment with CASIN. (B)-(C) Accelerating Rotarod was used to evaluate motor balance and coordination across S1/S2 mouse cohorts. The performance of expressing and non-expressing animals with and without CASIN treatment was compared and the latency to fall (in s) was measured. Performance was assessed at (B) 11-16 M: significant differences were found between synON16M+CASIN vs synOFF16M+CASIN (*), synOFF16M+CASIN vs synON16M (***), synON16M+CASIN vs synON16M (***) and synOFF16M vs synON16M (***), and (C) 19-24 M: significant differences occurred between synON24M vs synOFF24M (***), synON24M vs synON24M+CASIN (**), synON24M vs synOFF24M+CASIN (***), synOFF24M vs synON24M+CASIN (***) and synON24M+CASIN vs synOFF24M+CASIN (***) (n = 12 male and female mice, repeated-measures two-way ANOVA, *p<0.05, **p<0.01, ***p<0.001; data: mean ± SEM). (D)-(E) Gaussia luciferase activity analysis was performed on SEC fractions from S1/S2 mice aged (D) 16 M and (E) 24 M. Comparisons included groups with and without S1/S2 expression, as well as with or without CASIN treatment (n = 6 male and female mice per group, data: mean ± SEM). (F) Quantification of the AUC for total luciferase activity in SEC fractions. (G) Quantification of S1/S2 transgene expression in lysates from S1/S2 mice performed by capillary-based Western blot analysis (Simple Western), normalized to Actin.

To further uncover molecular mechanisms driving PD pathogenesis upon aging, we performed snRNA-seq on micro-dissected brains, using the 10X platform (**Fig.4 A**). To this end, we isolated nuclei from brain slices of mice from 6, 16, and 24 M old mice (synON6M, synOFF6M, synON16M, synlateON16M, synOFF16M, SynON24M, SynlateON24, SynOFF24M, n = 27 mice) with 2-3 replicate libraries from four animals for each genotype and condition. We obtained 105,689 high quality nuclei for analysis (**Suppl.Fig.1**). Unbiased clustering and cell type annotation based on known cell type-specific markers and marker genes from mousebrain.org (Zeisel et al. 2018) (**Suppl. Table 1**) identified 35 distinct cell types, including 12 inhibitory GABA neuronal clusters and 13 excitatory Vglut1 neuronal subclusters. Besides neuronal cell clusters we also identified six non-neuronal clusters including Astrocytes (ASC), Oligodendrocytes (OLG), Oligodendrocyte progenitor cells (OPC), Microglia (MGL), Choroid Plexus (CHOR), and Vascular cells (VSC) (**Fig.4 B-D**).

**Figure 4:**
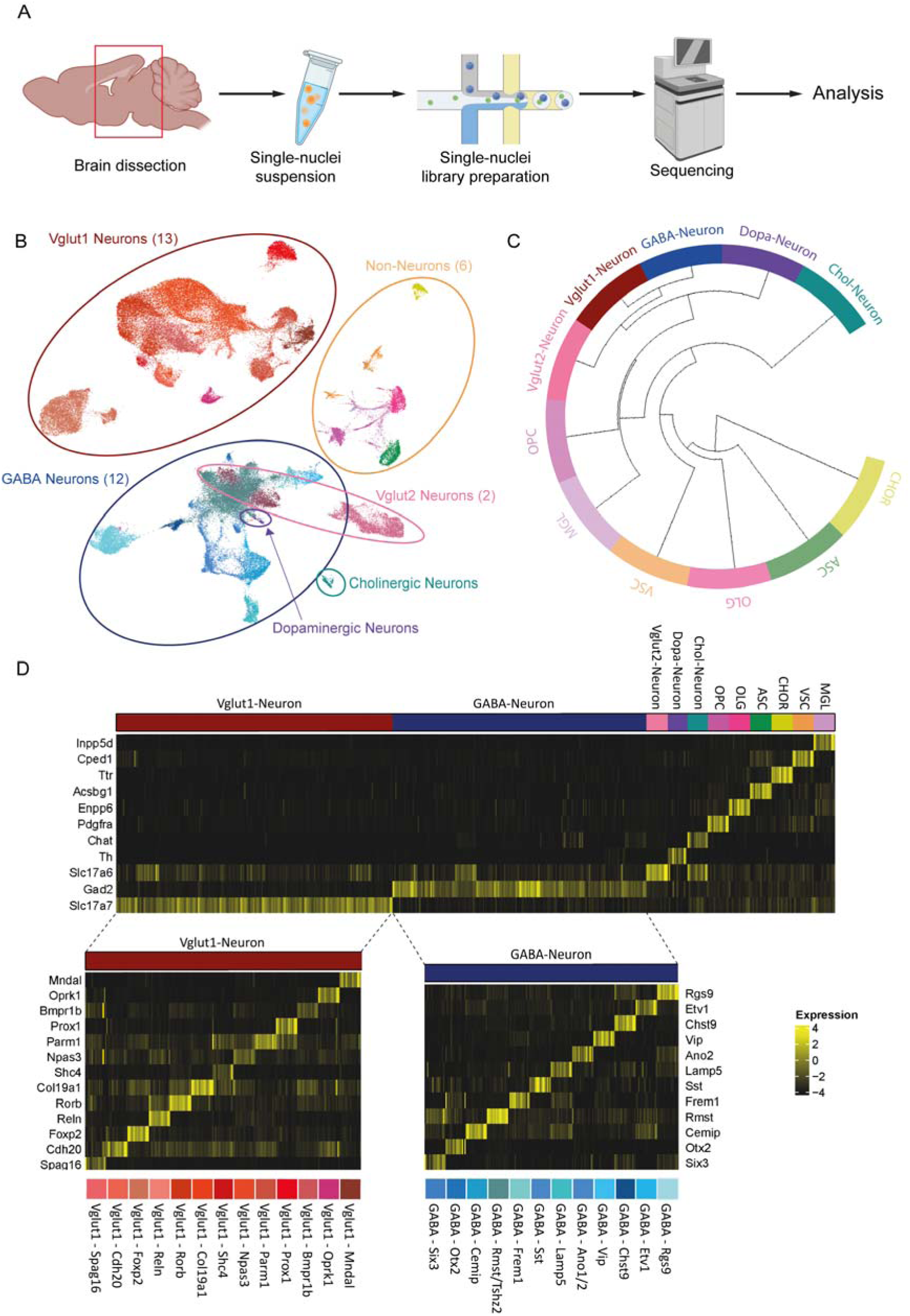
(A) Schematic workflow of the snRNA-seq approach. Nuclei were isolated from S1/S2 PD mice and used to generate gene expression libraries. Those were sequenced and proceeded to analysis (n = 4 animals per condition). Created with Biorender. (B) UMAP visualization of unbiased clustering with 105,689 quality-controlled cells. In total, 35 distinct cell types could be annotated and (C) hierarchical clustering of the main cell types. Distances between non-neuronal and neuronal clusters were the largest. (D) Canonical marker genes and marker genes from mousebrain.org showcased clear differential expression patterns between clusters.

To characterize the impact of the presence of S1/S2 oligomers on the distinct cell types, we assessed the relative abundance of S1/S2 transcripts across every cell type under all transgene expressing conditions. We found that the neuronal cell type GABA-Rgs9 exhibited the highest S1/S2 expression among all annotated cell types (**Fig.5 A**). Notably, the GABA-Rgs9 cluster represents GABAergic Basal Ganglia Neurons (BGNs) based on transcriptional data markers from mousebrain.org (**Suppl. Table 1**). Basal ganglia is a brain region crucial for motor control and central for motor impairments in PD. GABAergic neurons are important for movement regulating by forming inhibitory pathways within the basal ganglia. Additionally, GABA-Rgs9 cluster shows a high number of differentially expressed genes (DEGs) in PD mice (**Suppl.Fig.2**), which implied a strong differential response of these neurons to the presence of the oligomers. We therefore focused our further analysis on the GABA-Rgs9 cluster.

**Figure 5:**
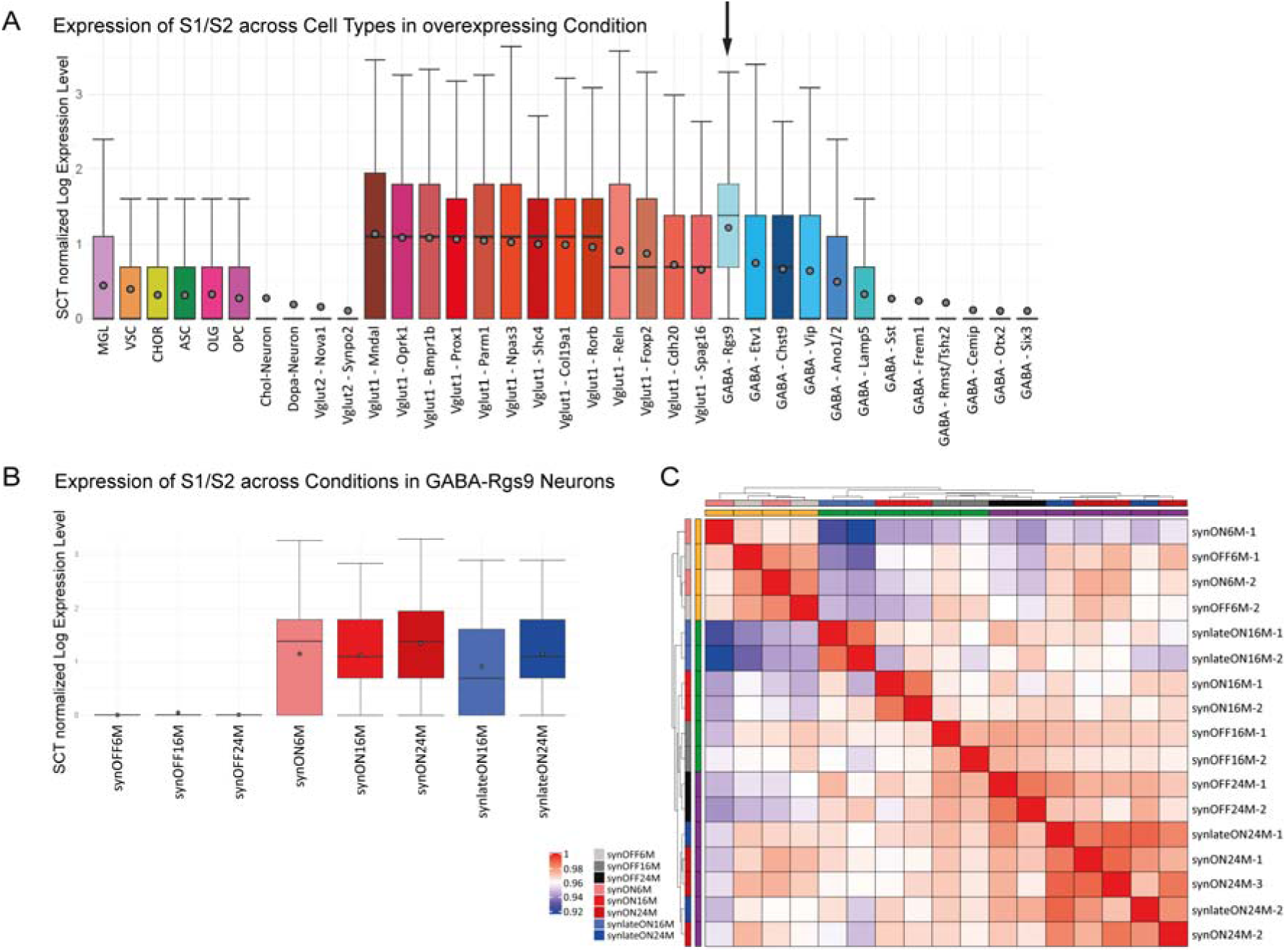
(A) Expression of S1/S2 transcript in S1/S2 expressing conditions across annotated cell types. GABA-Rgs9 neurons showed the highest overall expression of S1/S2. GABA-Rgs9 neurons also represent GABAergic BGNs. (B) S1/S2 expression in GABA-Rgs9 neurons were comparable between the S1/S2 expressing conditions, while the synOFF conditions showcased near-zero expression levels. (C) Similarity heatmap of all samples in GABA-Rgs9. (Euclidean distance with Complete-linkage clustering). At 16 M all samples were most similar to their replicates but at 24 M the synON and synlateON replicates formed a cluster together and at 6 M even the synON and synOFF clustered together. Boxplots with interquartile range (whiskers) and mean average (dots).

Within the GABA-Rgs9 cluster, we next examined the level of S1/S2 expression per cell across all conditions. We also performed a similarity analysis using all 11,706 detected genes in the GABA-Rgs9. The levels of S1/S2 transcripts were consistent among the transgene-expressing conditions, whereas the OFF conditions displayed, as expected, near-zero S1/S2 expression levels (**Fig.5 B**). A similarity heatmap based on the expression of all genes revealed sub clustering of synON24M and synlateON24M replicates at 24 M, a pattern not observed at 16 M. At 6 M, the synON and synOFF replicates clustered together which also corresponds to their similar motor performance at this age (**Fig.5 C**), as 16 M old S1/S2 mice displayed only minor motor deficits.

To delineate likely underlying molecular mechanisms of the motoric phenotype, DEGs were identified in GABA-Rgs9 cells by employing sample-wise pseudo-bulking for GABA-Rgs9 cells and performing Wald testing via the DESeq2 package (Love, Huber, and Anders 2014). Given that motor deficits were present in synlateON and synON animals at both 16 and 24 M compared to controls, we focused on DEGs present in synON vs synOFF and synlateON vs synOFF comparisons that are also shared between the two timepoints. These genes were compared to the DEGs from the comparison of synON6M vs synOFF6M, as there were no motoric deficits in synON6M mice. We derived two central gene signatures from this comparison: i) the “PD Beginning” signature of genes that are differentially expressed as response to S1/S2 expression prior to symptom and remain differentially expressed during motor deficits (DEGs present at both 6 M and in the intersection of 16 and 24 M), and ii) the "PD Signature" that represents genes differentially expressed only upon severe motoric PD (DEGs present at the intersection of 16 and 24 M that are not present at 6 M, **Fig.6 A**). The PD Signature comprised 102 genes, the PD Beginning 8 genes (**Suppl. Table 3**). Additionally, using bulk Mass Spectrometry (MassSpec), we obtained a protein PD Signature based on differential comparisons that are identical to the comparisons for obtaining the RNA PD Signature (**Suppl.Fig.3, Suppl. Table 4**). A final PD network was constructed by combining the snRNA-seq PD Signature, the snRNA-seq PD Beginning signature, and the bulk MassSpec protein PD Signature into a single representative network showing the most central genes of the signatures (**Fig.6 B**). The snRNA-seq signature networks were based on synON24M transcriptional data in the GABA-Rgs9 cell type.

**Figure 6:**
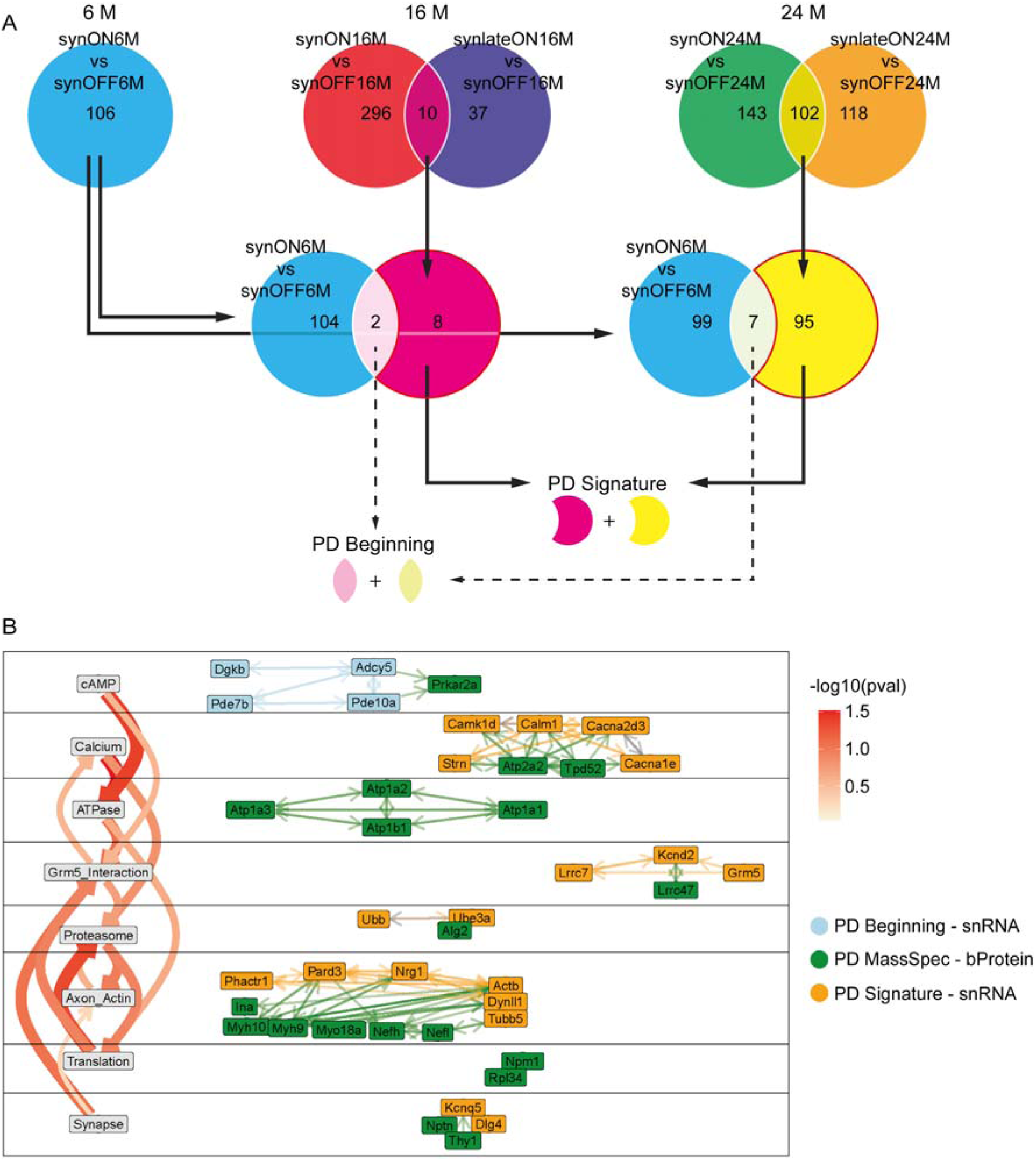
(A) Defining the “PD Signature” and “PD Beginning” gene set. Based on the rotarod data we decided to only consider intersection genes between the comparisons “synON vs synOFF” and “synlateON vs synOFF” at 16 M and 24 M for PD Signatures. From those intersections the genes that were not differentially expressed in “synON6M vs synOFF6M” were assigned to “PD Signature” while genes present in “synON6M vs synOFF6M” and the intersections were defined as “PD Beginning”. (B) Functional network using the central regulators from snRNA-seq PD Signature, PD Beginning genes, and bulk Protein (bProtein) PD Signature from MassSpec data. The interaction between genes/proteins and the p-values (pval) for topic-topic interactions were obtained based on omniPath and STRING databases. Network only showed topics with at least two genes/proteins and the top 2 topic-topic connections for each topic. The network suggests an upstream role of cAMP even before onset of motor deficits. This is followed by Calcium pathway and Grm5. Grm5 acts as an intersection between the more upstream pathways and the downstream effectors like Proteasome or Axon/Actin. Especially Axon/Actin showed high number of strongly interconnected genes/proteins. Additionally, it should be noted that the "Synapse" topic serves as a fallback category for genes/proteins with broad synaptic functions. Most genes/proteins assigned to other topics are also located in the synapse but have more specific subcellular roles.

The functional module “cAMP” included all dysregulated “PD Beginning” (blue) genes, including downregulation of cAMP homeostasis genes (*Adcy5, Pde7b, Pde10a*) and reduced protein levels of the cAMP target Prkar2a/Protein kinase A (PKA) in all S1/S2 expressing PD conditions. This suggests dyshomeostasis of cAMP-levels and reduced PKA activity to be an early impairment in the course of the disease and to be directly associated to elevated levels of S1/S2 oligomers. The calcium pathway containing downregulation of calcium influx channel gene expression (*Cacna2d3, Cacna1e*) alongside an upregulation of calcium binding signaling genes (*Calm1, Calm3*; **Suppl. Table 3**) was identified as a response to S1/S2 oligomers in response to cAMP signaling, likely as an adaptive response to reduced Calcium signaling. Both "cAMP" and "calcium" exerted strong direct and indirect effects on the glutamate metabotropic receptor 5 “Grm5_Interaction". Genes upstream (*Lrrc7*) and downstream (*Kcnd2*) of the glutamate metabotropic receptor 5 (*Grm5*), as well as *Grm5* itself, were downregulated in the PD Signature. Interestingly, *Grm5* Interaction appears to function as a connector, as it receives signals from the upstream modules cAMP and Calcium, but also from the downstream modules Translation and Synapse, while at the same time effecting the downstream modules of Proteasome and Axon/Actin. The Proteasome and Axon/Actin modules exhibited the strongest ingoing connections, indicating more downstream roles during the course of the disease. Within the Axon/Actin module (Actin: *Actb, Nrg1, Pard3, Phactr1*, Myosin/Dynein: MYH10, MYH9, MYO18A, *Dynll1*, Tubulin: *Tubb5*, NEFH, NEFL) the expression levels of actin genes were decreased, while Myosin/Dynein/Tubulin protein levels were increased, implying a significant change in axonal transport mechanisms (**Fig.6 B, Suppl. Table 2,4**). Notably, CDC42 activity can be a major regulator of cytoskeletal organization and synaptic activity (Nunes, Madeira, and Fonseca 2024).

The data predicts proteasomal dysfunction to be a central and downstream mechanism for motor dysfunction in the PD model (**Fig.6 B**). To test this *in-silico* findings experimentally, we determined protein degradation mechanisms in full-brain lysates. We investigated both, the ubiquitin-proteasome system (UPS) and the autophagic-lysosomal pathway, which have also been previously shown to be dysregulated in PD patients (Lim and Tan 2007; Lynch-Day et al. 2012). While the autophagy system (**Suppl.Fig.5 A-C**) and the Chymotrypsin- and Trypsin-enzymatic protease activity (**Suppl.Fig.5 G-H**) was not affected, reduced Caspase-like enzymatic protease activity confirmed downregulation of the UPS in synlateON24M or synON24M animals compared to the control mice (synOFF24M) (**Fig.7 A-C**). This is in line with our *in-silico* results, which obtained UPS associated genes (*Ubb, Ube3a*) as central genes of our PD Signature network (**Fig.6 B**).

**Figure 7:**
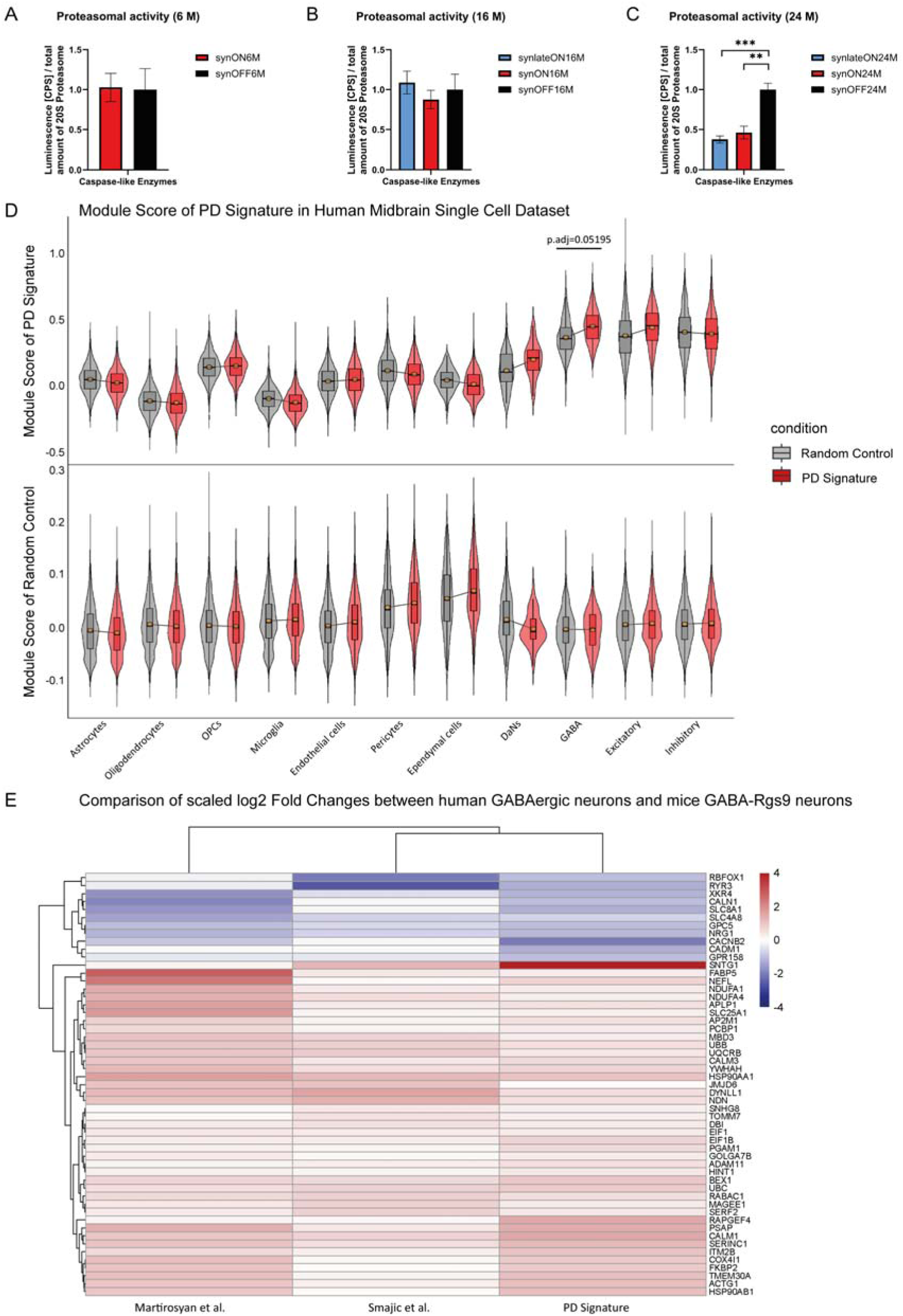
(A-C) Determination of proteasomal activity of Caspase-like enzymes of full-brain lysates from S1/S2 animals by proteasome activity assay. Comparison of short-, long- and non-expressing animals with (A) 6 M, (B) 16 M and (C) 24 M of age. Luminescence signal (counts per second, CPS) is proportional to proteasomal activity and was normalized to total amount of 20S proteasome determined by Western blot analysis (student’s t-test, ***<0.001, **<0.01; data: mean ± SEM). (D) Top 20 PD Signature genes and 20 randomly sampled genes were used to calculate a Module Score on the human midbrain snRNA-seq dataset Smajic et al., 2022. P values were calculated by Wilcoxon test on sample wise pseudo-bulked samples. Reflecting our finding in mice GABA-Rgs9 Neurons (GABAergic BGNs), the PD Signature is found to be most affecting the GABAergic Neurons in the human midbrain. This effect is not observed in the random control. (E) Heatmap of scaled log2 Fold Changes of genes from the PD Signature that were concordantly changed across mice GABA-Rgs9 synON24M vs synOFF24M, Smajic et al., human PD vs Control in GABAergic neurons, and Martirosyan et al. PD vs Control in GABAergic neurons. We obtained a total of 53 concordant genes, including genes we deemed as central to our PD Signature, such as Ubb, Dynll1, or Calm1.

Finally, we correlated our PD Signature to a snRNA-seq dataset of human PD midbrains (Smajić et al. 2022). Integrating our top 20 central PD Signature genes with the human dataset, there was a substantial increase in the PD vs control module scores in Dopaminergic Neurons (DaN), GABAergic Neurons and Excitatory Neurons (**Fig.7 D**), which was not observed in a random control experiment (**Fig.7 D**). Additionally, we used another snRNA-seq dataset of human PD midbrains with 15 PD patient samples and 14 controls (Martirosyan et al. 2024). We performed pseudo-bulked DGE analysis in GABAergic neurons for both datasets (Suppl. Table 6, 7) and compared the log2 fold changes from our PD Signature from mice GABA-Rgs9 neurons to the log2 fold changes in the human GABAergic neurons (Fig. 7 E, Suppl. Fig. 6). We identified 53 genes with concordant expression changes in both human and mouse GABAergic neurons (Fig. 7 E). These 53 genes included the central PD Signature genes such as *Ubb*, *Calm2*, or *Nrg1* (Fig. 7 E, Suppl. Fig. 6). This data suggests that the PD Signature is also differentially affected in human PD.

Our findings suggest that early PD involves disrupted cAMP and calcium signaling, reducing ATP production and impairing Grm5 function. This contributes to proteasomal and axon/actin system dysfunction in GABA-Rgs9 neurons (mice) and other neurons (humans), driving motor deficits in aging-related PD.

Aging is the causative driver for the onset of PD in the α-syn model. We therefore further identified an Aging Signature (similar to the identification of the PD Signature) by comparing the DGE synOFF16M vs synOFF6M” (6 M ➔ 16 M aging), the synOFF24M vs synOFF16M (16 M ➔ 24 M), and the synOFF24M vs synOFF6M” (6 M ➔ 24 M). Aging Signature genes were identified as genes that were concordantly changed throughout all aging stages, meaning only genes exhibiting the same direction of DE across all the three comparisons have been considered associated with aging and were included in the Aging Signature (**Fig.8 A**). As previously conducted for the PD Signature, the Aging Signature derived from MassSpec data was also integrated into the Aging network (**Suppl.Fig.3, Suppl. Table 5**). The functional network for aging was constructed using the same methodology as for the PD Signature (**Suppl.Fig.2, Suppl.Fig.5**). We grouped genes related to mitochondrial function, due to their abundance, under the category "Mitochondria" encompassing all mitochondria related genes (Methods Inference of Central regulators, **Fig.8 B, Suppl.Fig.4**, **Suppl. Table 5**). Since the Aging Signature network contained nearly twice as many regulators compared to the PD Signature network, our analyses now comprised the top 40 non-mitochondrial central regulators.

**Figure 8:**
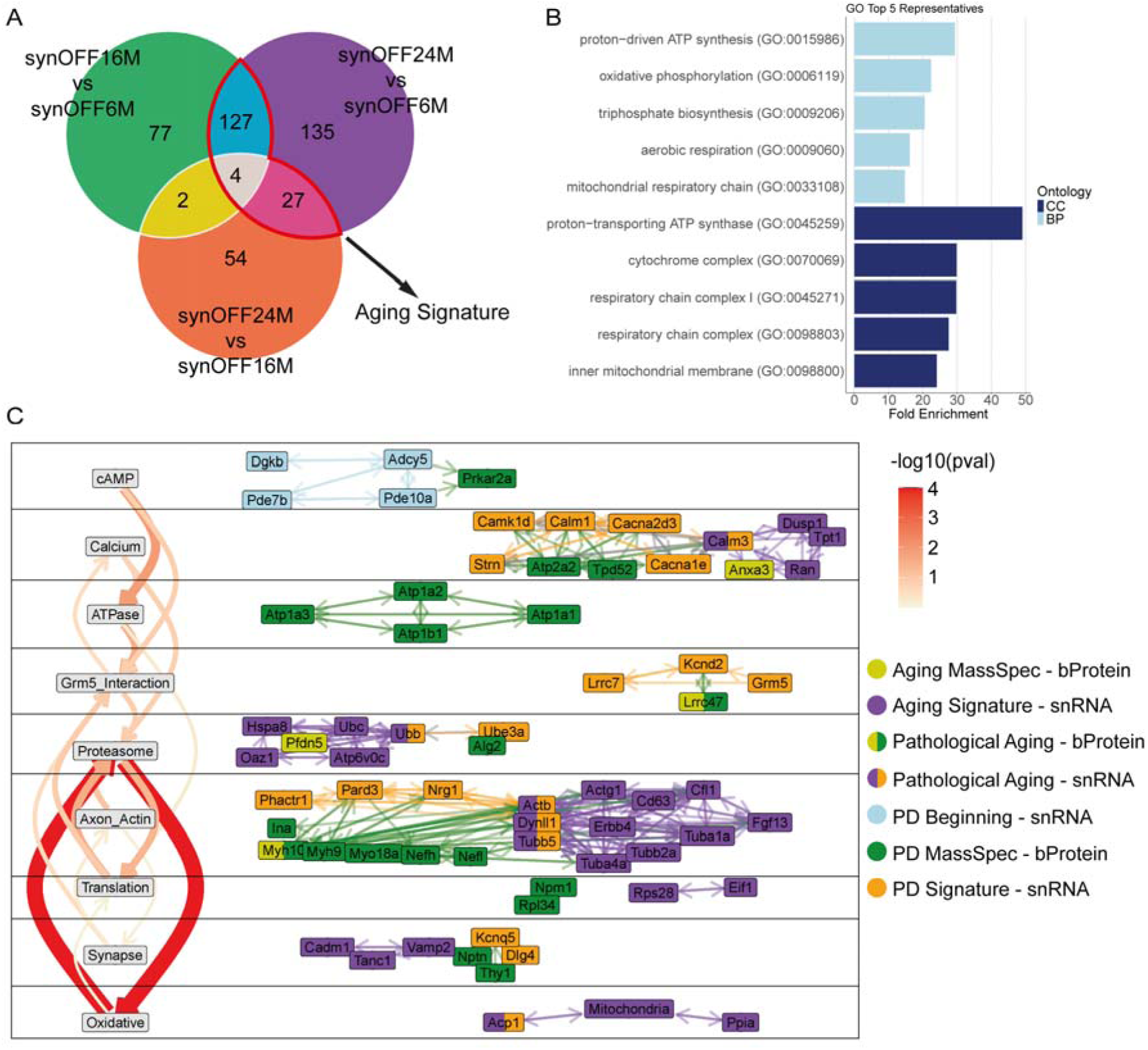
(A) Aging Signature contains all genes differentially expressed in “synOFF24M vs synOFF6M” as well as in “synOFF16M vs synOFF6M” or “synOFF24M vs synOFF16M”. (B) The Aging Signature was heavily influenced by genes specific to mitochondrial function and obscured other topics. Thus, all those genes were summarized under the term “Mitochondria”. (C) Final functional network combining the PD network and the top 40 central regulators from the non-mitochondrial Aging Signature. Notably, aging affected many same functional topics found in PD but also introduced new strong topic-topic interaction pathways, especially the interaction between Proteasome and Oxidative damage. We also found six genes that play a central role in both PD and aging and thus were defined as “Pathological Aging - snRNA”: Calm3, Ubb, Actb, Dynll1, Tubb5, Acp1 and two “Pathological Aging – bProteins”: Lrrc47 and Myh10.

The Aging Signature held functional modules that were very similar to the ones already identified in the PD Signature: Calcium, Proteasome, Axon/Actin, and Oxidative Damage. Interestingly, there was a direct overlap including eight genes/proteins (*Calm3, Ubb, Actb, Dynll1, Tubb5, Acp1*, MYH10, LRRC47), that we designated as "Pathological Aging” signature. These genes/proteins likely serve as central regulators in both PD and aging (**Fig.8 C)**. Expression of the Pathological Aging signature genes were also consistently upregulated in the human snRNA-seq PD datasets (**Suppl.Fig.6**), implying a role for these genes also in the aging human brain in regards to PD. The data further implies that dyshomeostasis of cAMP-levels and reduced PKA activity are early impairments specific to the presence of S1/S2 oligomers, since dyshomeostasis of cAMP-levels and reduced PKA activity were not present in the Aging Signature.

Together, we suggest that α-syn oligomers evoke dyshomeostasis of cAMP-levels and reduced PKA activity as early response and are necessary to develop PD symptoms, however, the presence of these oligomers alone is not sufficient for disease development. For PD disease development molecular mechanisms of aging (cytoskeleton rearrangement (axon-actin), oxidative damage and proteasomal dysfunction) are necessary and sufficient for the development of the phenotypic disease.

## Discussion

The findings presented here provide significant insights into the mechanistic interplay between aging-related processes and PD progression in a model driven by α-syn oligomer formation. We demonstrate that the levels of aggregated α-syn do not show significant differences across the different age groups for the life-long α-syn expression. Our data suggests that the onset of motor symptoms is not directly correlated to α-syn oligomer load and α-syn oligomer formation precedes motor decline in aged mice. Only aging in combination with α-syn oligomers determines the PD motor phenotype. Support for the idea that aging rather a direct effect of α-syn oligomers is the determinant for pathological α-syn propagation and clinical symptoms in PD comes from the study by Vena den Berge et al. (Van Den Berge et al. 2021). Here intragastrical injection of α-syn preformed fibrils into young rats did not result in CNS pathology, whereas old rats clearly showed α-syn pathology throughout the brain. This indicates that aging influences the brain’s susceptibility to α-syn-induced pathology rather than the aggregation process per se.

Further evidence also comes from our snRNA seq data. SnRNA-seq identified that the PD Signature is indeed partially overlapping with an Aging Signature in GABA-Rgs9 that are most affected by the expression of the α-syn oligomers. Attenuation of Cdc42 activity, which is known to restore the aging system to a more youthful level, clearly alleviated PD symptoms in aged mice without reducing α-syn oligomers, demonstrating that aging per se is necessary and sufficient to drive PD onset and progression in this model.

Aging remains the single largest risk factor for most chronic diseases. It is well known that aging is the main risk factor for neurodegeneration (Hou et al. 2019) and also for the development of PD. But why is aging necessary that a neurodegenerative disease like PD develops? In geroscience studies it is investigated how cellular and molecular changes associated with aging contribute to the onset and progression of these diseases. Our α-syn inducing PD model is a strong experimental validation of the concept of geroscience: Α-syn oligomers in this mouse model evoke dyshomeostasis of cAMP-levels and reduced PKA activity as early response and are necessary to develop PD symptoms. But the oligomers and their mode of action is not sufficient for phenoconversion. The α-syn oligomer load does not correlate with disease symptoms per se. For PD disease development molecular mechanisms of aging like cytoskeleton rearrangements, oxidative damage, and proteasomal dysfunction are necessary. As strong proof of this concept we demonstrate that pharmacological attenuation of elevated activity of CDC42 in brain cells upon aging (L. Wang et al. 2007) ameliorates motor dysfunction in aged PD mice, but without affecting α-syn oligomer levels.

The attenuation of CDC42 activity rejuvenates fundamental aging processes, such as cytoskeletal organization and synaptic activity. Our data suggest that α-syn oligomers reduce PKA signaling by disrupting cAMP homeostasis (e.g., via downregulation of *Adcy5* and *Pde10a*). Since PKA inhibits CDC42 activity, reduced PKA signaling might further exacerbate CDC42 activation. Elevated CDC42 activity, already linked to aging-related cellular changes (Umbayev et al. 2023), may reach a pathological threshold when this aging-associated increase combines with α-syn oligomers reduce cAMP/PKA signaling and thus the inhibiting function of PKA towards CDC42 is impaired. We have already demonstrated that α-syn oligomers bind specifically CDC42 effector proteins and likely by these means are also able to influence CDC42 signaling (Schnack et al. 2008). This could explain why PD symptoms manifest during aging and aging is necessary for symptom onset, as aging amplifies CDC42 dysregulation.

Our snRNA-seq and network analyses identified key pathways dysregulated by aging, including cAMP and calcium signaling, proteasomal function, and cytoskeletal dynamics. The aging signature shared substantial overlap with PD-associated pathways, particularly in proteasomal and axon/actin modules, implicating also on the level of transcripts the shared mechanism of "pathological aging” as a driving force for phenoconversion of PD. Interestingly, the cAMP pathway was already deregulated before symptoms onset and remained deregulated over time, implying that this dysregulation might be necessary, but not sufficient for causing symptoms, and that only the combined presence of both cAMP and aging-related calcium signaling deficits results in PD phenotypes. Calm3, critical member of our Pathological Aging gene set, encodes calmodulin, a critical calcium-binding protein and a key calcium sensor, that regulates synaptic activity (Aravind et al. 2019). Calcium imbalance increases with aging and is thought to play a key role in PD (Calì, Ottolini, and Brini 2014). We find Calm1 and Calm3 to be upregulated in human PD brain. PD patients’ brains show elevated intracellular calcium, especially in substantia nigra neurons, where L-type Cav1.3 calcium channels are highly expressed (C. S. Chan, Gertler, and Surmeier 2009; Hurley et al. 2013). The calcium channel blocker isradipine improved motor function and reduced neurodegeneration in animal models (Q.-M. Wang et al. 2017). Finally, excess calcium promotes α-syn aggregation by exposing its aggregation-prone NAC region (Lautenschläger et al. 2018). Our findings of enrichment of α-syn oligomers in aged PD animals suggest a feedback loop where excess calcium drives α-syn aggregation to further negatively influence PKA/CDC42 signaling.

It is well known that proteasomal dysfunction is linked to aging but also to PD, with atypical ubiquitination of proteins like α-syn promoting insoluble aggregates (Buneeva and Medvedev 2022; Moon et al. 2020). Loss of proteostasis is also considered a hallmark of aging (López-Otín et al. 2013). Our PD Signature shows changes in *Ubb* and *Ube3a*, while we also found altered levels of *Ubb* in our aging signature, supporting a role for proteasomal dysfunction as part of pathological aging and PD. Consistent with snRNA-seq data, caspase-like proteasome activity is significantly reduced in 24-month-old animals, supporting proteasomal dysfunction as a late-stage effect of α-syn oligomers (Lindersson et al. 2004; McKinnon et al. 2020). Among the pathological aging genes we also found Acp1, which has been previously linked to oxidative damage during aging and age related diseases (Matos et al. 2018). Acp1 might therefore form a link between aging, PD and proteasome dysfunction by oxidative damage. Direct evidence though of an ACP1-CDC42 interaction is lacking, but evidence suggests ACP1 influences CDC42 via upstream regulators (Nakano and Mabuchi 2006; Zou et al. 2024; Hu et al. 2011).

In summary, we report that α-syn oligomers reduce activity in the cAMP/PKA pathway in an early non-symptomatic phase while molecular mechanisms of aging like calcium signaling dysregulation, cytoskeletal rearrangements, oxidative damage and proteasomal dysfunction are likely major causative driver for the aging-associated onset of PD.

---

***Suppl. Fig. 1:*** *(A) Scatterplot showcasing the number of Unique Molecular Identifiers (x-axis, nUMIs) and the number of genes (y-axis, nGene) per barcode across different conditions. Filtering thresholds are marked at 2500 for nUMI and 1500 for nGene on the respective axes. The color of the points represents the percentage of mitochondrial genes detected per barcode, with darker points indicating higher values. (B) Similar to (A), but here the color of the points represents the percentage of ribosomal genes. The scatterplots indicate comparable sequencing across conditions and barcodes with high mitochondrial or ribosomal gene percentages are found mostly in the filtered barcodes. (C) Results after data integration, showing that all clusters and cell types are represented across all conditions, even though cell numbers do vary between conditions: synOFF6M 23,324 - synON6M 24,015 - synOFF16M 5,346 - synlateON16M 3,407 - synON16M 5,453 - synOFF24M 5,921 - synlateON24M 15,016 - synON24M 23,207. (D) The nUMI was comparable between all conditions*.

***Suppl. Fig. 2:*** *(A) Pipeline for differential expression gene (DEG) analysis. (B) To verify that batch effects do not significantly impact the results, p-value histograms were generated for comparisons between Batch1 vs. Batch2 and synOFF24M vs. synOFF6M, where the batch was a confounding factor. The histograms show minimal batch effects. (C) Scatterplot depicting the total number of DEGs in each cell type for synON vs synOFF comparisons, plotted against the number of UMIs (nUMI) per cell type with a red regression line and the 95% confidence interval (grey zone). Cell types with low nUMI scatter further from the expected regression line. This is likely due to insufficient nUMI and thus differential hits are more likely technical differences. Notably, the GABA-Rgs9 Neurons, our cell type of interest, shows a significantly higher number of differential hits compared to other cell types. (D) Network inference pipeline is illustrated. Using the single nucleus count matrix of one condition for the signature genes in PIDC, an adjacency matrix with confidence scores for all interactions is inferred. The top interconnected genes are then integrated with curated interaction databases, OmniPath and STRING, to construct a biologically relevant network. Central regulators are identified via hypergeometric testing and manually annotated into functional topics. This network is then combined with bulk-protein data and other signatures. Topic-topic interaction scores are calculated to generate the final network graph*.

***Suppl. Fig. 3:*** *(A) Analysis Pipeline for bulk Protein data from MassSpec. It consists out of filtering the raw MaxLFQ counts, followed by normalization and imputation of missing values. (B) Most proteins were detected across all 32 replicates. (C) However, the two replicates from Batch2 showed a lower detection rate compared to those from Batch1. To address this, proteins only detected in both batches in at least two samples were kept. Subsequent filtering was performed to retain proteins measured in at least 75% of replicates in a minimum of one condition. (D) After filtering and normalization, the remaining missing values were imputed, which did not alter the dataset’s structure. (E) By incorporating the batch effect as a covariate in the differential analysis design, this method effectively removed the batch effect from the dataset*.

***Suppl. Fig. 4:*** *(A) Top 5 KEGG-Terms representing the genes in the Aging Signature. Oxidative Phosphorylation, which represents the electron transport system in mitochondria, showcases highly significant Fold Enrichment compared to other terms. (B-C) This mitochondrial bias is also evident when examining the top central regulators from the Aging Signature network. Most of the top 40 regulators are genes specific to mitochondrial function. (C) By summarizing and excluding mitochondrial-specific genes, we identify the relevant top 40 central regulators. As done for the PD Signature network, "No Info" vertices are removed, and connections to regulators are replaced with direct connections between the regulators. This network was further annotated, and inter-topic edges was removed for clearer visualization. (D) Gene Ontology (GO) and KEGG terms for the genes excluding mitochondrial-specific ones predominantly include metabolic and aging-related terms*.

***Suppl. Fig. 5:*** *(A-F) Determination of autophagy of full-brain lysates from S1/S2 animals by analyzing of autophagic marker LC3. Comparison of short-, long- and non-expressing animals with (A) 6 M, (B) 16 M and (C) 24 M of age. (D-E) Representative western blot pictures with autophagic marker LC3 and GAPDH as loading control. (G-H) Determination of proteasomal activity of Caspase-, Chymotrypsin- and Trypsin-like enzymes of full-brain lysates from all cohorts by proteasome activity assay. Comparison of the animals with (G) 16 M and (H) 24 M of age. Luminescence signal (counts per second, CPS) is proportional to proteasomal activity and was normalized to total amount of 20S proteasome determined by Western blot analysis (student’s t-test, ***<0.001, **<0.01; data: mean ± SEM)*.

***Suppl. Fig. 6:*** *(A) Scatter plot showing the Log2 fold changes for the PD Signature genes. Y-axis represents synON24M vs synOFF24M comparison in the GABA-Rgs9 cell type in our dataset while the x-axis represents PD vs Control comparison in GABAergic Neurons in the human snRNA dataset of human midbrain samples for Smajic et al. (Smajic et al., 2022) and Martirosyan et al. (Martirosyan et al. 2024). We observe a concordant change between our and the human dataset for all six Pathological Aging genes (Acp1, Actb, Calm3, Ubb, Dynll1, Tubb5). Ubb, Dynll1, and Calm3 were concordant in both human datasets, while Acp1, Actb and Tubb5 were only upregulated in one of the human datasets*.

## Methods

### Generation and Housing of Transgenic Animals

Generation and a detailed description of the phenotype characterization has been previously published (Kiechle et al. 2019). The animals were housed at the Animal Research Center, Ulm University under standardized conditions. As the S1/S2 model is a Tet-off system, the transgene expression was suppressed by adding 100 mg/ml doxycycline in the drinking water, which was additionally sweetened with 10 g/l glucose. All animals were housed in groups in open polycarbonate type II long cages under controlled room temperature, humidity, a 12 h dark/light cycle and food and water *ad libitum*. The cages were enriched with nesting material and polycarbonate houses. Each subgroup of the experimental cohorts included 12 animals (balanced mixed sex groups) which underwent behavioral testing once a week.

### Behavioral Experiment – Accelerating Rotarod

Accelerating Rotarod (five lane Rotarod, Med Associates) was used to determine motor balance and coordination of all S1/S2 mouse cohorts, comparing animals with different ages and length of S1/S2 expression. For each time point, every mouse had to undergo three consecutive trials with breaks of 5 min in between. For one trial, the rotation of the rod increased from 4 to 40 rpm over a period of 300 s and the latency to fall was recorded by a software (Rotarod 1.2.0 software, Med Associates). For the analysis, mean values were used.

### Protein Complementation Assay

The S1/S2 mouse model includes a bioluminescent protein complementation assay (PCA) expressing human wildtype α-syn fused to either the N- or C-terminal part of a Gaussia luciferase. Those halves are called S1 and S2, and are expressed simultaneously. After the aggregation of the expressed α-syn, the halves form a functional active luciferase that can be detected by the administration of the substrate coelenterazine (Danzer et al. 2011). The expression takes place under the neuron-specific CamKIIα promoter (Mayford et al. 1996) and is inducible employing a Tet-off system driven by Doxycycline (Kiechle et al. 2019). The drinking water of the breeding cages always included doxycycline to avoid any transgene expression during embryonic development.

### CASIN Treatment

CASIN (Xcess Biosciences M60040) was freshly prepared on a weekly basis prior to each round of injections. Initially, the substance was dissolved in dimethyl sulfoxide (DMSO) to achieve a 100 mM stock solution, which was subsequently diluted using a (2-Hydroxypropyl)-beta-cyclodextrin solution (Sigma H5784). Mice received intraperitoneal injections at a dose of 25 mg/kg once daily, around midday, for four consecutive days. This cycle was followed by a one-week break, after which the four-day injection regimen was repeated. Treatments commenced when the mice were either 12 or 20 M old. Control animals received injections containing only the vehicle in equivalent volumes. CASIN was supplied as a lyophilized powder, and a single batch was consistently used throughout the entire experimental period.

### Perfusion

Transcardial perfusion with 1x PBS of the whole animal was performed to rapidly remove the blood from the animals. At the age of 6, 16 or 24 M, the mice were terminally anesthetized with ketamine (100 mg/kg) and xylazine (16 mg/kg). After testing reflexes, the skin of the animal was cut ventrally under the ribs cage. The ribs were cut on the left side and lifted to expose the heart. The needle attached to a syringe filled with 50 ml PBS was put into the left ventricle and the right atrium was cut immediately to allow the blood and fluid to flow out of the organism. Afterwards, the brain was removed from the skull and fresh frozen in liquid nitrogen.

### Protein Extraction of Mouse Brain Tissue

For full-brain lysate preparation, one hemisphere per brain was homogenized in PBS with a ratio of 1 g tissue per 10 ml PBS. After homogenization using a TissueLyser II (Qiagen) for 2 x 2 min at 25 Hz, the homogenates were centrifuged at 20,800 g for 30 min at 4 °C. Then protein concentration of the supernatant was determined by BCA assay (Thermo Fisher Scientific) and used for the further experiments (SEC, Western blot analysis, Simple Western experiments and Proteasome Activity assay).

### BCA Assay

The protein concentrations of the full-brain lysates were determined by colorimetric two-component Pierce BCA Protein Assay Kit (Thermo Fisher Scientific) like indicated in the manufacturer’s instructions. The lysates were diluted 1:5, 1:10 and 1:20, and PBS was used as blank. The concentrations of the used protein standards (bovine serum albumin) ranged from 0-2 mg/ml (0, 125, 250, 500, 750, 1000, 1500, 2000 µg/ml). After mixing the fluorescent dye with the standard samples, lysates and blank, everything was incubated at 37 °C for 30 min and the fluorescence intensity was measured in duplicates using a Microplate reader (SPECTROstar Nano) at wavelength 562 nm.

### Size-Exclusion Chromatography

The full-brain lysates were used for SEC performed with the Superdex 200 Increase 10/300 GL column (Cytiva). The column was connected to an Äkta pure system (Cytiva) and got equilibrated with two column volumes (CV) of filtered (0.22 µm filters) and degased PBS before use. A 1 ml loop was used for automated sample injection and 600 µl of each lysate was loaded onto the column. A flow rate of 0.75 ml/min for PBS was used for elution of the proteins from the column. The maximum pressure was set to 5 MPa and the eluted proteins were monitored by UV absorbance at 280 and 215 nm. The eluate was collected in fractions of 500 µl in deep-well plates for later experiments. 200 µl were used for duplicate measurement of luciferase activity and 100 µl were used for dot blot analysis. Determination of the molecular mass of the eluted proteins was performed using the Gel Filtration Calibration Kit (Cytiva) with the following standard proteins: Conalbumin (75 kDa), Ovalbumin (44 kDa), Carbonic anhydrase (29 kDa), Ribonuclease A (13.7 kDa) and Aprotinin (6.5 kDa). The void volume (v_o_) was determined using Blue Dextran.

### Human Gaussia Luciferase Assay

Duplicates of 100 µl of each protein-containing fraction of SEC were used for measurement of luciferase activity. Aggregated α-syn with luciferase halves reconstitutes the whole Gaussia luciferase which oxidatively decarboxylates the substrate coelenterazine with emission of light as the result. Coelenterazine (P.J.K) stock was prepared previously to the experiments to a final concentration of 1 mg/ml in methanol and stored at -80 °C. The same batch was used for all measurements. The working solution of coelenterazine was prepared in Opti-MEM (40 µM) and incubated at RT for 25 min without light. 100 µl of the cell permeable substrate was directly added to each sample before measurement by the automated dispenser module of the plate reader (Victor X3 microplate reader, Perkin-Elmer). Luminescence was then measured at 480 nm with a signal integration time of 1 s. Luminescence signal was afterwards normalized to protein concentration. Lysates from animals without S1/S2 expression which did not undergo SEC but were also tested by luciferase activity assay to verify that there is no remaining luciferase expression.

### Dot Blot

Additionally, the SEC fractions were tested for total human α-syn signal by using a Dot blot apparatus (Minifold I, Whatman). The system was used as described by the manufacturer. 100 µl of each protein-containing SEC fraction was filtered through a nitrocellulose membrane by vacuum to transfer the proteins. After drying the membrane for 10 min at RT, it was blocked with 1x Roti-Block (Carl Roth) for 1 h at RT. It was then incubated overnight with primary antibody in 1x Roti-Block (anti-α-syn 15G7 antibody, 1:100) at 4 °C. The membrane was washed 3x 10 min with PBS-T and then incubated with secondary antibody in 1x Roti-Block (anti-rat HRP-conjugated goat antibody, 1:5000) for 2 h at RT. After additional 3 washing steps with 10 min with PBS-T each. Images were taken with the Fusion SoloS Imager (Vilber) with HRP substrate (SuperSignal West Pico PLUS, Thermo Fisher Scientific). The quantification of α-syn signal of SEC fractions was performed using the Fusion software.

### Simple Western

The Jess system (Bio-techne) was used to conduct a capillary-based Western blot analysis (Simple Western). The full-brain lysates underwent dilution and preparation according to the manufacturer’s protocol. Briefly, the samples were mixed with Simple Western Sample Buffer to achieve a final concentration of 0.4 mg/ml, followed by denaturation at 95 °C for 5 minutes. The samples, along with primary (anti-α-syn Syn1 antibody, BD, 1:100) and secondary (anti-mouse HRP antibody) antibodies, chemiluminescent substrate and wash buffer were dispensed into the 384-well plate. The Jess system executes all assay steps automatically by placing Simple Western assay buffers, capillaries and the prepared assay plate. Protein separation occurs within the capillaries as they migrate the stacking and separation matrix, leading to the immobilization of separated proteins on the capillary wall. Afterwards, target proteins are identified by primary antibody and immunodetection takes place by a HRP-conjugated secondary antibody and chemiluminescent substrate. The Compass software (Bio-techne) determines automatically molecular weight and AUC for immunodetected proteins. The expression of Syn-1 was normalized to beta-actin.

### Western Blot

For Western blotting, 37 µl of each sample was used, including 25 µg protein of the full-brain lysates and 14 µl β-mercaptoethanol, and got heated for 5 min at 95 °C. The samples were separated by sodium dodecyl sulfate-polyacrylamide gel electrophoresis (SDS-PAGE), using polyacrylamide gels with a 16 % separating and a 5 % stacking gel in a XCell SureLock Mini-Cell Electrophoresis system (marker: PageRuler Prestained protein ladder, Thermo Fisher Scientific). The gels were started with 80 V and were increased to 100 V when the samples reached the separating gel (running buffer: 25 mM Tris, 192 mM Glycin, 1 g/l SDS). Next, the proteins were transferred from the gel to a pre-activated PVDF membrane using a XCell SureLock XCell II Blot Module (Thermo Fisher Scientific) with 25 V for 2 h (blotting buffer: 25 mM Tris, 192 mM Glycin, 20 % Methanol). The transfer was verified by PonceauS staining. The following step included blocking of the membrane with TBS-T buffer (10 mM Tris-HCl, 150 mM NaCl, 0.05 % Tween-20) containing 5 % milk powder for 1 h. Overnight incubation of blots with primary antibodies (anti-LC3, Novus biologicals) diluted at 1:1000 in blocking solution occurred at 4 °C. The membranes underwent three 10-minute washes with 1x TBS-T to eliminate unbound primary antibodies. The next step involved incubation with secondary antibody (anti-rabbit IgG, diluted at 1:5,000 in blocking solution) for 2 h at RT, utilizing HRP-conjugated goat secondary antibodies (Promega). Following three additional 10-minute washes with 1x TBS-T, images were captured using a Fusion SoloS Imager (Vilber) with HRP substrate (SuperSignal West Pico PLUS, Thermo Fisher Scientific). Densiometric analysis was performed using the Fusion software.

For Western blots with the antibody against the 20S proteasome α1/α2/α3/α5/α6/α7 subunits (Santa Cruz), only 12 % polyacrylamide gels and 40 µg protein per sample was used. Here, a nitrocellulose membrane was used instead of PVDF and the blocking solution was 1x Roti-Block in water (Carl Roth).

### Proteasome Activity Assay

To evaluate proteasomal activity in mouse full-brain lysates a modified version of a commercially available indirect enzyme-based luminescent assay (Promega, G8621), utilizing a substrate designed for caspase-like activity was employed. The setup was adapted to measure proteasomal activity in total protein extracts from brain tissue according to Strucksberg et al. in 2009 (Strucksberg et al. 2010). The full-brain lysates were diluted to a concentration of 0.2 mg/ml with PBS. Subsequently, 50 µl, equivalent to 10 µg of total protein, was mixed with 50 µl of the luminescent reagent containing Ultra-Glo Luciferase and the signal peptide coupled to luciferin. All samples were incubated for 1 h in a 96-well plate at RT and the luminescence was then measured with a signal integration time of 1 s (Victor X3 Microplate reader, Perkin-Elmer). The luminescent signal strength in each examined lysate is directly proportional to the overall peptidase activity, representing the summation of proteasomal activity and non-specific peptidase activities originating from other enzymes present in the protein extract. The approach involved measurement with and without the addition of 30 µM of the irreversible and specific proteasomal inhibitor adamantine-acetyl-(6-aminohexanoyl)3-(leucinyl)3-vinyl-(methyl)-sulfone (AdaAhx3L3VS, Calbiochem). Proteasomal activity was determined as value by subtracting the non-specific background activity from the total peptidase activity. For a conclusive determination of specific proteasomal activity correlated with the proteasome quantity, the calculated proteasomal activity value underwent normalization using the densiometry data obtained from the 20S proteasome α1/α2/α3/α5/α6/α7 subunits immunoblot analysis.

### Nuclei Isolation

For nuclei isolation from mouse brain tissue, around 100 mg of brain tissue was cut from one hemisphere including 0 mm Bregma to -5 mm Bregma using a mouse coronal brain matrix (Cell Point Scientific). Two of those brain pieces from two different mice were pooled for the extraction. All procedures were carried out on ice. 1400 µl of homogenization buffer (320 mM Sucrose, 5 mM Cacl_2_, 3 mM Mg(Ac)_2_, 10 mM Tris HCl pH 8, 0.1 mM EDTA pH 8, 0.1 % NP-40, 1 mM β-mercaptoethanol, 0.4 U/µl RiboLock in H_2_O) was added to a douncer including the brain tissue and mechanical disruption with around 40 strokes was performed using pestle A followed by pestle B. The resulting homogenate was then passed through a 70 µm Flowmi cell strainer, followed by a 40 µm Flowmi filter. Next, 700 µl of the homogenized suspension was mixed with 450 µl of the working solution (50 % Opti-Prep, 5 mM CaCl_2_, 3 mM Mg(Ac)_2_, 10 mM Tris HCl pH 8, 0.1 mM EDTA, 1 mM β-mercaptoethanol in H_2_O). A gradient was prepared using the following components: 300 µl of 40 % Opti-Prep (40 % Opti-Prep, 96 mM Sucrose, 5 mM CaCl_2_, 3 mM Mg(Ac)_2_, 10 mM Tris HCl pH 8, 0.1 mM EDTA, 0.03 % NP-40, 0.12 U/µl RiboLock in H_2_O), 750 µl of 30 % Opti-Prep (30 % Opti-Prep, 134 mM Sucrose, 5 mM CaCl_2_, 3 mM Mg(Ac)_2_, 10 mM Tris HCl pH 8, 0.1 mM EDTA pH 8, 1 mM β-mercaptoethanol, 0.04 % NP-40, 0.17 U/µl RiboLock in H_2_O) and 700 µl of the mixture of homogenized tissue in working solution. The gradient underwent centrifugation at 10,000 g for 5 min at 4 °C. Following centrifugation, approximately 200 µl of the nuclei were aspirated and transferred into a 1.5 ml low-DNA-binding tube. To this, 250 µl of 2 % BSA and 0.12 U/µl RiboLock in PBS was added. Subsequent centrifugation at 2,000 g for 3 min at 4 °C was performed and the supernatant was discarded. The resulting pellet was resuspended in 250 µl of 2 % BSA and 0.12 U/µl RiboLock, followed by a second round of centrifugation at 2,000 g for 3 min at 4 °C. After discarding the supernatant, the pellet was once again resuspended in 250 µl of 2 % BSA and 0.12 U/µl RiboLock and filtered through a 40 µm Flowmi cell strainer into a low-DNA-binding tube. Further centrifugation at 2,000 g for 3 min at 4 °C was carried out. The pellet obtained was then resuspended in 50 µl of 1x nuclei buffer (1x nuclei buffer of 10x Genomics, with 1 mM DTT, 1 U/µl RiboLock in H_2_O). For counting nuclei and quality check for nuclear membrane integrity, nuclei were stained with DAPI. The final nuclei were then directly used for the Single-nuclei RNA isolation protocol.

### Single-nuclei RNA Sequencing

The Chromium 3’ Single Cell Library Kit (10x Genomics) was used to generate the Gel Beads-In-Emulsion (GEMs), following the guidelines provided by the manufacturer. Shortly, 22,000 of the isolated nuclei from mouse brain tissue were introduced into barcoded Gel Beads using the Chromium Controller. Following GEM-RT incubation, cDNA samples underwent recovery, purification and amplification through a cDNA amplification reaction. Quality assessments on the amplified cDNA were conducted using a High Sensitivity DNA Kit (Agilent) on a TapeStation platform. Libraries were subsequently generated through the processes of fragmentation and adaptor ligation. Sample Index PCR was executed and the resulting purified libraries underwent assessment on TapeStation using a High Sensitivity DNA Kit to evaluate fragment quality. The single-nucleus libraries were then sent to Novogene (United Kingdom) for sequencing with the NovaSeq 6000.

### Sequencing and alignment of single nucleus samples

Paired 150bp snRNA-seq was performed using the 10X Genomics Gene Expression (GEX) 3′protocol with a NovaSeq 6000 sequencer. For the alignment of reads, a custom reference was created by adding the sequences of the S1/S2 transgene and the CamkIIa promoter to the mm10 mouse reference genome. Count matrices were obtained using the *cellranger count 7.1* (Zheng et al. 2017) pipeline, including introns. Six samples were mapped using the bwUni2.0 High-Performance Computing infrastructure.

### Quality Control and Integration of cells and counts

The unfiltered count matrices were loaded into R and corrected for ambient mRNA using *SoupX 1.6.0* (Young and Behjati 2020) with default settings, adjusting “tfidfMin” settings between 0.9 and 1.3 depending on the sample. Seurat objects were created for each sample and subsequently merged. Cells were filtered out based on the following criteria: number of unique molecular identifiers (nUMI) < 2500, number of genes (nGene) < 1500, mitochondrial gene percentage > 3%, ribosomal gene percentage > 1.5%, or log10(Genes/nUMI) < 0.85. Next, doublets were removed using *scDblFinder 1.16.0* (Germain et al. 2021) with default values. In addition, doublets based on sex were removed using *cellXY 0.99.0* with default values Normalization was performed using the SCTransform function on 4000 variable features with glmGamPoi method implemented in *Seurat 5.0.1* (Hao et al. 2024), and top 50 embeddings were obtained via *scVI (scvi-tools 1.1.1)* (Gayoso et al. 2022) integration for sex, age, batch, and number of pooled animals. Clustering was done using the Leiden algorithm and visualized with Uniform Manifold Approximation and Projection algorithm (UMAP). Clusters represented by few samples, less than 100 cells, or a single batch, and not of conditions of interest, were removed. Clusters driven by ribosomal or mitochondrial genes, as well as markers of hindbrain and olfactory cell types, were also discarded. The steps from normalization onward were repeated until no further clusters needed removal. Final integration was performed using *harmony 1.2.0* (Korsunsky et al. 2019) with an integration diversity penalty (theta) of 2, followed by final clustering based on the top 30 harmony components and UMAP visualization. Each subsequent clustering for annotation of sub-cell types was computed following the same procedure.

### Annotation of Cell types

Clusters were annotated in a hierarchical manner using literature, the Mouse Brain Atlas (mousebrain.org) (Zeisel et al. 2018) and markers identified via the FindConservedMarkers function in Seurat. First, neurons and non-neuronal cells were distinguished using mainly canonical markers, such as but not limited to Rbfox3 (neurons), Mbp (oligodendrocytes), Acsbg1 (astrocytes), Pdgfra (oligodendrocyte precursor cells), Cx3cr1 (microglia), Colec12 (vascular cells), and Ttr (choroid plexus cells). Neurons were further classified into Vglut1, Vglut2, GABA, cholinergic, and dopaminergic neurons. Vglut1 and GABA neurons were further sub-annotated. A full list of markers and cell types can be found in the Suppl. Table 1.

### Differential Gene Expression Analysis for snRNA data

DEGs were identified using the Wald test from the DESeq2 package (Love, Huber, and Anders 2014). Cells were pseudo-bulked per sample for each annotated cell type, normalized, and all ribosomal and mitochondrial-DNA genes were excluded. All genes that were not expressed in at least three samples at an expression level of three were excluded in the analysis. To ensure comparability, cell number variability between batches and conditions were limited to a maximum of 1.5-fold difference via random subsampling. Covariates for the design were obtained via the sva package (Leek et al. 2012) and p-values were recalculated using *fdrtool 1.2.18* (Strimmer 2008). Multiple testing correction was applied using the Bonferroni-Holm method. A cutoff of 0.05 for adjusted p-values for significance was used. Potential remaining batch effects were controlled by differential analysis between the batches and any significant “batch-genes” were removed from comparisons with confounding designs.

### Defining Gene Signatures

The Aging Signature was defined as follows: All DEGs from the “synOFF24M vs synOFF6M” comparison that were also concordantly significant in the “synOFF16M vs synOFF6M” or “synOFF24M vs synOFF16M” comparisons. Genes that were only significant in “synOFF16M vs synOFF6M” and “synOFF24M vs synOFF16M” were not included into the Aging Signature as those genes do not change concordantly through the ages.

The PD Signature was defined as follows: We focused on differential genes present in both the “synON vs synOFF” and “synlateON vs synOFF” comparisons at 16 M. The same approach was applied for 24 M. These intersection genes between synON and synlateON were then compared to DEGs from the “synON6M vs synOFF6M” comparison. From this comparison, we created two signatures: Genes present at both 6 M and in the intersection were designated as “PD Beginning.” Additionally, the "PD Signature" was defined as all intersection genes not present at 6 M. The results were combined to obtain the final “PD Signature” and “PD Beginning” gene sets.

### Inference of Networks

Using PIDC, the most interconnected genes were chosen based on T. E. Chan et al. (T. E. Chan, Stumpf, and Babtie 2017). PIDC, based on Partial Information Decomposition (PID), infers statistical dependencies between triplets of genes from a single-cell gene expression matrix by assigning a confidence score to each interaction. For network inference, synON24M GABA-Rgs9 neuron counts were used for the PD Signature and synOFF24M GABA-Rgs9 neuron was used for the Aging Signature. Only connections with a top 10 % confidence score were used to build an interaction network using curated public databases of known interactions. The OmniPath database (Türei, Korcsmáros, and Saez-Rodriguez 2016) was accessed using OmnipathR 3.11.16 (Türei et al. 2021) and combined with interactions having a confidence score above 0.600 from STRING version 12.0.0 (Szklarczyk et al. 2019). The final network was constructed by retrieving the interactions from the databases between the PIDC genes, allowing for a maximum of one interconnecting gene.

### Inference of central Regulators

Each vertex/gene was tested for significant over-connectivity using hypergeometric testing, comparing the number of connections of the gene in the obtained graph to the number of connections of the gene in the database. Vertices were ranked according to their adjusted p-values, and the top 20 genes were designated as central regulators for the PD Signature.

For the Aging Signature, mitochondria genes were biased to be defined as central regulator due to the high interconnectivity between genes specific to mitochondrial function, which would mask possible other effects. To find other relevant aging genes besides mitochondria function we summarized all genes represented in the KEGG term “Oxidative Phosphorylation” or in any GO-Cellular Component term containing “mitochondrial,” “respiratory chain,” “respirasome,” “NADH,” or “oxidoreductase” as “Mitochondria”. From the remaining genes, the top 40 genes were identified as central regulators as the number of significantly central genes were close to double in the Aging Signature compared to the PD Signature.

### Topic categorization and interaction between topics

Central regulators were categorized into functional topics using the databases of GO (Aleksander et al. 2023), KEGG (Kanehisa et al. 2023), UniProt (The UniProt Consortium 2021), and GeneCard (Safran et al. 2021). An interaction score between categories was calculated as follows: A distribution of connections was synthesized by permuting the topic assignment of genes 10,000 times and recording the number of interactions between the topics. Using this distribution, p-values were obtained and -log10 of the p-values served as the interaction scores for the observed number of connections between the topics. The top two interaction partners for each topic were considered for further analysis. For final visualization, only topics with at least two assigned genes/proteins were included.

### Label-free proteomic data synthesis of mouse brain tissue

Tissue lysates containing 100 µg total protein (16-32 µL) were filled to 40 µL with 100 mM triethylammonium bicarbonate (TEAB) and Tris(2-carboxyethyl)phosphine hydrochloride (TCEP, final concentration 5 mM) and 2-chloroacetamide (final concentration 10 mM) were added. Samples were incubated at 95 °C for 10 min. After cool down, 180 µL 100 mM TEAB and a trypsin/LysC (Promega) solution (protein-to-enzyme ratio 50:1,) were added and samples incubated at 37 °C overnight. The digestion was stopped by addition of 50 µL 6 % TFA (final 1 %). Peptides were fractionated using STAGE Tips (AttractSPE Disks Bio SDB, Affinisep) into 3 fractions using 15 % acetonitrile (ACN, fraction 1), 24 % ACN (fraction 2) and 70 % ACN in 20 mM NH4-Formiat (pH10). The fractions were vacuum dried and resuspended in 12 µL 0.5 % TFA. Peptides were separated using an UltiMate 3000 RSLCnano system and a PepMap100 C18, 20 × 0.075 mm, 3 μm trap column (Thermo) and a PepMap100 C18, 50 × 0.050 mm, 2 μm analytical column (Thermo). Mobile phase of the loading pump (trap column) was 0.05 % TFA/2 % MeOH (flow rate: 5 µL/min), the mobile phase of the nano pump (analytical column) was A: 4 % DMSO/0.1 % formic acid and B: 4 %DMSO/76 % acetonitrile/0.1 % formic acid (flow rate: 150 nL/min) and peptides were separated with a 3 h gradient. Peptides were infused into a QExactive mass spectrometer (Thermo) and measured with data-dependent acquisition (Top15). Proteins were identified using MaxQuant 2.0.3.0 and the Mus musculus reference proteome from UniProt (downloaded 3rd February 2022). A FDR of 1 % was used for peptide and protein identification and protein quantification was performed with the MaxLFQ algorithm (Cox, Hein et al. 2014).

### Mass Spectrometry Data Analysis

MassSpec data included four replicates per condition from two batches. First, proteins detected in at least two samples in both batches were retained. Subsequently, the data was further filtered for proteins measured in a minimum of 75% of replicates for at least one condition. All methods thereafter, including Normalization, missing value imputation, and differential expression analysis, were performed using the DEP package and its wrapper functions. Furthermore, normalization between batches was facilitated by certain samples being purposefully remeasured in both batches.

For imputation of the remaining missing values, protein expression values were classified as either missing at random (MAR) or missing not at random (MNAR). A value was defined as MNAR if the proteins was detected in at least 75% of replicates of one condition while the other condition had 75% missing values. The imputation algorithms were chosen in the DEP function "impute" with "knn" implementation set for MAR values and "MinProb" set for MNAR values. A more detailed explanation of the algorithms can be found in the official DEP package documentation. Three replicates (synlateON16-4, synON16-3, and synOFF16-4) made a subcluster independent of their conditions. Thus, differential analysis was performed without those three samples. Differential expression analysis was conducted with the wrapper function for limma in DEP, using sex and batch information as covariates, and significance set at a Bonferroni-Holm adjusted p-value below 0.1.

### Comparing Results with Human data

To validate our findings against published human data, we utilized the dataset from Smajic et al. (2022) (Smajić et al. 2022). The “CADPS2+” cell type was excluded from the analysis. We computed a Module Score using the top 20 PD signature genes + Calm3, employing the AddModuleScore function of Seurat. As a random control, we subsampled an equivalent number of genes as for the PD signature from the human dataset and calculated a corresponding Module Score. P values between conditions were calculated by unpaired Wilcoxon testing on sample-wise pseudo-bulked samples. Additionally, we compared the log2FoldChanges between the synON24M versus synOFF24M conditions in mouse GABA-Rgs9 neurons and the PD versus Control conditions in human GABAergic neurons.

Additionally, we used another snRNA-seq dataset of human PD midbrains with 15 PD patient samples and 14 controls (Martirosyan et al. 2024). Cells were filtered and processed according to the original publication. A cluster was annotated as GABAergic Neurons when all four relevant markers showed enriched expression (*GAD1*, *GAD2*, *SNAP25*, *SYT1*). We performed pseudo-bulked DGE analysis using covariates from sva and sex as known covariate in GABAergic neurons for both datasets and compared the log2 fold changes from our PD signature from mice GABA-Rgs9 neurons to the log2 fold changes in the human GABAergic neurons. For this, all log2 Fold Changes were scaled to -4 to 4 to make datasets comparable.

### Quantification and Statistical Analysis

Statistical analysis was performed and graphs were created with Prism (GraphPad Prism 10). For behavioral analysis, repeated-measures two-way ANOVA was performed. All tests for significance were two tailed with a=0.05. Values are presented as mean ± SEM (standard error of the mean). P-values <0.05 were considered as statistically significant with significance levels *p<0.05, **p<0.01 and ***p<0.001.

## Data and code availability

The raw sequencing data that was generated in this study is available from ArrayExpress with the identifier (to be followed). Detailed metadata, the processed single-cell data and all other source data are available on Zenodo (to be followed). The code generated for the analysis and the code used to generate the figures is available on GitHub: https://github.com/DanzerLab/snRNA_PDMouseModel_Age/

## Acknowledgements

We thank R. Bück, S. Meier, A. Erk and A. Jesionek for their excellent work and technical support.

## Author Contribution

V. Bopp, M. Kiechle and K. M. Danzer designed the project. V. Bopp performed experiments and analyzed the data. J. Kühlwein supported snRNA-seq experiments. J. LeeBae performed snRNA-seq and MassSpec analysis. P. Öckl performed MassSpec experiments and analysis. J. Kühlwein, V. Grozdanov, B. Mayer, B. Möhrle and H. Geiger provided intellectual input. V. Bopp, J. LeeBae and K. M. Danzer wrote the manuscript. All authors reviewed and approved the manuscript.

## Abbreviations

α-syn: α-synuclein
Adcy: adenylyl cyclase
ASC: astrocytes
AUC: area under the curve
BGNs: basal ganglia neurons
CASIN: CDC42 activity specific inhibitor
CDC42: cell division control protein 42 homolog
CHOR: choroid plexus
CPS: counts per second
DaN: dopaminergic neurons
DEGs: differentially expressed genes
GEMs: gel beads-in-emulsion
Grm5: glutamate metabotropic receptor 5
M: month
MAR: missing at random
MassSpec: Mass Spectrometry
MGL: microglia
MNAR: missing not at random
nUMI: number of unique molecular identifiers
OLG: oligodendrocytes
OPC: oligodendrocyte progenitor cells
PCA: protein complementation assay
PD: Parkinson’s Disease
PDE: phosphodiesterase
PID: partial information decomposition
PKA: protein kinase A
SEC: Size-Exclusion Chromatography
snRNA-seq: single-nuclei RNA-sequencing
UMAP: Uniform Manifold Approximation and Projection algorithm
UPS: ubiquitin-proteasome system
V_o_: void volume
VSC: vascular cells

## References

Aleksander, Suzi A, James Balhoff, Seth Carbon, J Michael Cherry, Harold J Drabkin, Dustin Ebert, Marc Feuermann, et al. 2023. ‘The Gene Ontology Knowledgebase in 2023’. Genetics 224 (1): iyad031. 10.1093/genetics/iyad031.

Aravind, P., Sarojini R. Bulbule, N. Hemalatha, R.L. Babu, and K.S. Devaraju. 2019. ‘Elevation of Gene Expression of Calcineurin, Calmodulin and Calsyntenin in Oxidative Stress Induced PC12 Cells’. Genes & Diseases 8 (1): 87–93. 10.1016/j.gendis.2019.09.001.

Buneeva, Olga, and Alexei Medvedev. 2022. ‘Atypical Ubiquitination and Parkinson’s Disease’. International Journal of Molecular Sciences 23 (7): 3705. 10.3390/ijms23073705.

Calì, Tito, Denis Ottolini, and Marisa Brini. 2014. ‘Calcium Signaling in Parkinson’s Disease’. Cell and Tissue Research 357 (2): 439–54. 10.1007/s00441-014-1866-0.

Chan, C. Savio, Tracy S. Gertler, and D. James Surmeier. 2009. ‘Calcium Homeostasis, Selective Vulnerability and Parkinson’s Disease’. Trends in Neurosciences 32 (5): 249–56. 10.1016/j.tins.2009.01.006.

Chan, Thalia E., Michael P.H. Stumpf, and Ann C. Babtie. 2017. ‘Gene Regulatory Network Inference from Single-Cell Data Using Multivariate Information Measures’. Cell Systems 5 (3): 251–267.e3. 10.1016/j.cels.2017.08.014.

Chu, Yaping, and Jeffrey H. Kordower. 2007. ‘Age-Associated Increases of α-Synuclein in Monkeys and Humans Are Associated with Nigrostriatal Dopamine Depletion: Is This the Target for Parkinson’s Disease?’ Neurobiology of Disease 25 (1): 134–49. 10.1016/j.nbd.2006.08.021.

Danzer, Karin M., Wolfgang P. Ruf, Preeti Putcha, Daniel Joyner, Tadafumi Hashimoto, Charles Glabe, Bradley T. Hyman, and Pamela J. McLean. 2011. ‘Heat-Shock Protein 70 Modulates Toxic Extracellular α-Synuclein Oligomers and Rescues Trans-Synaptic Toxicity’. FASEB Journal: Official Publication of the Federation of American Societies for Experimental Biology 25 (1): 326–36. 10.1096/fj.10-164624.

Florian, Maria Carolina, Karin Dörr, Anja Niebel, Deidre Daria, Hubert Schrezenmeier, Markus Rojewski, Marie-Dominique Filippi, et al. 2012. ‘Cdc42 Activity Regulates Hematopoietic Stem Cell Aging and Rejuvenation’. Cell Stem Cell 10 (5): 520–30. 10.1016/j.stem.2012.04.007.

Florian, Maria Carolina, Hanna Leins, Michael Gobs, Yang Han, Gina Marka, Karin Soller, Angelika Vollmer, et al. 2020. ‘Inhibition of Cdc42 Activity Extends Lifespan and Decreases Circulating Inflammatory Cytokines in Aged Female C57BL/6 Mice’. Aging Cell 19 (9): e13208. 10.1111/acel.13208.

Gayoso, Adam, Romain Lopez, Galen Xing, Pierre Boyeau, Valeh Valiollah Pour Amiri, Justin Hong, Katherine Wu, et al. 2022. ‘A Python Library for Probabilistic Analysis of Single-Cell Omics Data’. Nature Biotechnology 40 (2): 163–66. 10.1038/s41587-021-01206-w.

Germain, Pierre-Luc, Aaron Lun, Carlos Garcia Meixide, Will Macnair, and Mark D. Robinson. 2021. ‘Doublet Identification in Single-Cell Sequencing Data Using scDblFinder’. F1000Research 10:979. 10.12688/f1000research.73600.2.

Hao, Yuhan, Tim Stuart, Madeline H. Kowalski, Saket Choudhary, Paul Hoffman, Austin Hartman, Avi Srivastava, et al. 2024. ‘Dictionary Learning for Integrative, Multimodal and Scalable Single-Cell Analysis’. Nature Biotechnology 42 (2): 293–304. 10.1038/s41587-023-01767-y.

Hou, Yujun, Xiuli Dan, Mansi Babbar, Yong Wei, Steen G. Hasselbalch, Deborah L. Croteau, and Vilhelm A. Bohr. 2019. ‘Ageing as a Risk Factor for Neurodegenerative Disease’. Nature Reviews Neurology 15 (10): 565–81. 10.1038/s41582-019-0244-7.

Hu, Jinghui, Alka Mukhopadhyay, Peter Truesdell, Harish Chander, Utpal K. Mukhopadhyay, Alan S. Mak, and Andrew W. B. Craig. 2011. ‘Cdc42-Interacting Protein 4 Is a Src Substrate That Regulates Invadopodia and Invasiveness of Breast Tumors by Promoting MT1-MMP Endocytosis’. Journal of Cell Science 124 (Pt 10): 1739–51. 10.1242/jcs.078014.

Hurley, Michael J., Bianca Brandon, Steve M. Gentleman, and David T. Dexter. 2013. ‘Parkinson’s Disease Is Associated with Altered Expression of CaV1 Channels and Calcium-Binding Proteins’. Brain: A Journal of Neurology 136 (Pt 7): 2077–97. 10.1093/brain/awt134.

Ingelsson, Martin. 2016. ‘Alpha-Synuclein Oligomers-Neurotoxic Molecules in Parkinson’s Disease and Other Lewy Body Disorders’. Frontiers in Neuroscience 10:408. 10.3389/fnins.2016.00408.

Kamath, Tushar, Abdulraouf Abdulraouf, S. J. Burris, Jonah Langlieb, Vahid Gazestani, Naeem M. Nadaf, Karol Balderrama, Charles Vanderburg, and Evan Z. Macosko. 2022. ‘Single-Cell Genomic Profiling of Human Dopamine Neurons Identifies a Population That Selectively Degenerates in Parkinson’s Disease’. Nature Neuroscience 25 (5): 588–95. 10.1038/s41593-022-01061-1.

Kanehisa, Minoru, Miho Furumichi, Yoko Sato, Masayuki Kawashima, and Mari Ishiguro-Watanabe. 2023. ‘KEGG for Taxonomy-Based Analysis of Pathways and Genomes’. Nucleic Acids Research 51 (D1): D587–92. 10.1093/nar/gkac963.

Kiechle, Martin, Bjoern von Einem, Lennart Höfs, Patrizia Voehringer, Veselin Grozdanov, Daniel Markx, Rosanna Parlato, et al. 2019. ‘In Vivo Protein Complementation Demonstrates Presynaptic α-Synuclein Oligomerization and Age-Dependent Accumulation of 8-16-Mer Oligomer Species’. Cell Reports 29 (9): 2862–2874.e9. 10.1016/j.celrep.2019.10.089.

Korsunsky, Ilya, Nghia Millard, Jean Fan, Kamil Slowikowski, Fan Zhang, Kevin Wei, Yuriy Baglaenko, Michael Brenner, Po-ru Loh, and Soumya Raychaudhuri. 2019. ‘Fast, Sensitive and Accurate Integration of Single-Cell Data with Harmony’. Nature Methods 16 (12): 1289–96. 10.1038/s41592-019-0619-0.

Lautenschläger, Janin, Amberley D. Stephens, Giuliana Fusco, Florian Ströhl, Nathan Curry, Maria Zacharopoulou, Claire H. Michel, et al. 2018. ‘C-Terminal Calcium Binding of α-Synuclein Modulates Synaptic Vesicle Interaction’. Nature Communications 9 (1): 712. 10.1038/s41467-018-03111-4.

Leek, Jeffrey T., W. Evan Johnson, Hilary S. Parker, Andrew E. Jaffe, and John D. Storey. 2012. ‘The Sva Package for Removing Batch Effects and Other Unwanted Variation in High-Throughput Experiments’. Bioinformatics 28 (6): 882–83. 10.1093/bioinformatics/bts034.

Li, Jiahua, Ka Wai Ng, Chun Chau Sung, and Kenny K. K. Chung. 2024. ‘The Role of Age-Associated Alpha-Synuclein Aggregation in a Conditional Transgenic Mouse Model of Parkinson’s Disease: Implications for Lewy Body Formation’. Journal of Neurochemistry 168 (7): 1215–36. 10.1111/jnc.16122.

Lim, Kah-Leong, and Jeanne MM Tan. 2007. ‘Role of the Ubiquitin Proteasome System in Parkinson’s Disease’. BMC Biochemistry 8 (Suppl 1): S13. 10.1186/1471-2091-8-S1-S13.

Lindersson, Evo, Rasmus Beedholm, Peter Højrup, Torben Moos, WeiPing Gai, Klavs B. Hendil, and Poul H. Jensen. 2004. ‘Proteasomal Inhibition by α-Synuclein Filaments and Oligomers*’. Journal of Biological Chemistry 279 (13): 12924–34. 10.1074/jbc.M306390200.

López-Otín, Carlos, Maria A. Blasco, Linda Partridge, Manuel Serrano, and Guido Kroemer. 2013. ‘The Hallmarks of Aging’. Cell 153 (6): 1194–1217. 10.1016/j.cell.2013.05.039.

Love, Michael I., Wolfgang Huber, and Simon Anders. 2014. ‘Moderated Estimation of Fold Change and Dispersion for RNA-Seq Data with DESeq2’. Genome Biology 15 (12): 550. 10.1186/s13059-014-0550-8.

Lynch-Day, Melinda A., Kai Mao, Ke Wang, Mantong Zhao, and Daniel J. Klionsky. 2012. ‘The Role of Autophagy in Parkinson’s Disease’. Cold Spring Harbor Perspectives in Medicine 2 (4): a009357. 10.1101/cshperspect.a009357.

Martirosyan, Araks, Rizwan Ansari, Francisco Pestana, Katja Hebestreit, Hayk Gasparyan, Razmik Aleksanyan, Silvia Hnatova, et al. 2024. ‘Unravelling Cell Type-Specific Responses to Parkinson’s Disease at Single Cell Resolution’. Molecular Neurodegeneration 19 (1): 7. 10.1186/s13024-023-00699-0.

Matos, Andreia, Alda Pereira da Silva, Isanete Alonso, Maria João Marta, Filipa Albergaria, Ana Luiza Torres, Natércia Joaquim, Isabel Júlio, José Pereira Miguel, and Manuel Bicho. 2018. ‘P-382 - Oxidative Stress-Induced Biomarkers May Influence Lipid Profile in Young Adults’. Free Radical Biology and Medicine 120 (May):S161. 10.1016/j.freeradbiomed.2018.04.529.

Mayford, M., M. E. Bach, Y. Y. Huang, L. Wang, R. D. Hawkins, and E. R. Kandel. 1996. ‘Control of Memory Formation through Regulated Expression of a CaMKII Transgene’. Science (New York, N.Y.) 274 (5293): 1678–83. 10.1126/science.274.5293.1678.

McKinnon, Chris, Mitchell L. De Snoo, Elise Gondard, Clemens Neudorfer, Hien Chau, Sophie G. Ngana, Darren M. O’Hara, et al. 2020. ‘Early-Onset Impairment of the Ubiquitin-Proteasome System in Dopaminergic Neurons Caused by α-Synuclein’. Acta Neuropathologica Communications 8 (1): 17. 10.1186/s40478-020-0894-0.

Moon, Stuart P., Aaron T. Balana, Ana Galesic, Ananya Rakshit, and Matthew R. Pratt. 2020. ‘Ubiquitination Can Change the Structure of the α-Synuclein Amyloid Fiber in a Site Selective Fashion’. The Journal of Organic Chemistry 85 (3): 1548–55. 10.1021/acs.joc.9b02641.

Nakano, Kentaro, and Issei Mabuchi. 2006. ‘Actin-Capping Protein Is Involved in Controlling Organization of Actin Cytoskeleton Together with ADF/Cofilin, Profilin and F-Actin Crosslinking Proteins in Fission Yeast’. Genes to Cells: Devoted to Molecular & Cellular Mechanisms 11 (8): 893–905. 10.1111/j.1365-2443.2006.00987.x.

Nunes, Mariana, Natália Madeira, and Rosalina Fonseca. 2024. ‘Cdc42 Activation Is Necessary for Heterosynaptic Cooperation and Competition’. Molecular and Cellular Neurosciences 129 (June):103921. 10.1016/j.mcn.2024.103921.

Raket, Lars Lau, Daniel Oudin Åström, Jenny M. Norlin, Klas Kellerborg, Pablo Martinez-Martin, and Per Odin. 2022. ‘Impact of Age at Onset on Symptom Profiles, Treatment Characteristics and Health-Related Quality of Life in Parkinson’s Disease’. Scientific Reports 12 (January):526. 10.1038/s41598-021-04356-8.

Safran, Marilyn, Naomi Rosen, Michal Twik, Ruth BarShir, Tsippi Iny Stein, Dvir Dahary, Simon Fishilevich, and Doron Lancet. 2021. ‘The GeneCards Suite’. In Practical Guide to Life Science Databases, edited by Imad Abugessaisa and Takeya Kasukawa, 27–56. Singapore: Springer Nature. 10.1007/978-981-16-5812-9_2.

Schnack, C., K. M. Danzer, B. Hengerer, and F. Gillardon. 2008. ‘Protein Array Analysis of Oligomerization-Induced Changes in Alpha-Synuclein Protein-Protein Interactions Points to an Interference with Cdc42 Effector Proteins’. Neuroscience 154 (4): 1450–57. 10.1016/j.neuroscience.2008.02.049.

Silkis, I. 2001. ‘The Cortico-Basal Ganglia-Thalamocortical Circuit with Synaptic Plasticity. II. Mechanism of Synergistic Modulation of Thalamic Activity via the Direct and Indirect Pathways through the Basal Ganglia’. Bio Systems 59 (1): 7–14. 10.1016/s0303-2647(00)00135-0.

Skuladottir, Astros Th, Vinicius Tragante, Gardar Sveinbjornsson, Hannes Helgason, Arni Sturluson, Anna Bjornsdottir, Palmi Jonsson, et al. 2024. ‘Loss-of-Function Variants in ITSN1 Confer High Risk of Parkinson’s Disease’. Npj Parkinson’s Disease 10 (1): 1–4. 10.1038/s41531-024-00752-9.

Smajić, Semra, Cesar A. Prada-Medina, Zied Landoulsi, Jenny Ghelfi, Sylvie Delcambre, Carola Dietrich, Javier Jarazo, et al. 2022. ‘Single-Cell Sequencing of Human Midbrain Reveals Glial Activation and a Parkinson-Specific Neuronal State’. Brain: A Journal of Neurology 145 (3): 964–78. 10.1093/brain/awab446.

Strimmer, Korbinian. 2008. ‘Fdrtool: A Versatile R Package for Estimating Local and Tail Area-Based False Discovery Rates’. Bioinformatics 24 (12): 1461–62. 10.1093/bioinformatics/btn209.

Strucksberg, Karl-Heinz, Karthikeyan Tangavelou, Rolf Schröder, and Christoph S. Clemen. 2010. ‘Proteasomal Activity in Skeletal Muscle: A Matter of Assay Design, Muscle Type, and Age’. Analytical Biochemistry 399 (2): 225–29. 10.1016/j.ab.2009.12.026.

Szklarczyk, Damian, Annika L Gable, David Lyon, Alexander Junge, Stefan Wyder, Jaime Huerta-Cepas, Milan Simonovic, et al. 2019. ‘STRING V11: Protein–Protein Association Networks with Increased Coverage, Supporting Functional Discovery in Genome-Wide Experimental Datasets’. Nucleic Acids Research 47 (Database issue): D607–13. 10.1093/nar/gky1131.

The UniProt Consortium. 2021. ‘UniProt: The Universal Protein Knowledgebase in 2021’. Nucleic Acids Research 49 (D1): D480–89. 10.1093/nar/gkaa1100.

Türei, Dénes, Tamás Korcsmáros, and Julio Saez-Rodriguez. 2016. ‘OmniPath: Guidelines and Gateway for Literature-Curated Signaling Pathway Resources’. Nature Methods 13 (12): 966–67. 10.1038/nmeth.4077.

Türei, Dénes, Alberto Valdeolivas, Lejla Gul, Nicolàs Palacio-Escat, Michal Klein, Olga Ivanova, Márton Ölbei, et al. 2021. ‘Integrated Intra- and Intercellular Signaling Knowledge for Multicellular Omics Analysis’. Molecular Systems Biology 17 (3): e9923. 10.15252/msb.20209923.

Umbayev, Bauyrzhan, Yuliya Safarova Yantsen, Aislu Yermekova, Assem Nessipbekova, Aizhan Syzdykova, and Sholpan Askarova. 2023. ‘Role of a Small GTPase Cdc42 in Aging and Age-Related Diseases’. Biogerontology 24 (1): 27–46. 10.1007/s10522-022-10008-9.

Van Den Berge, Nathalie, Nelson Ferreira, Trine Werenberg Mikkelsen, Aage Kristian Olsen Alstrup, Gültekin Tamgüney, Páll Karlsson, Astrid Juhl Terkelsen, Jens Randel Nyengaard, Poul Henning Jensen, and Per Borghammer. 2021. ‘Ageing Promotes Pathological Alpha-Synuclein Propagation and Autonomic Dysfunction in Wild-Type Rats’. Brain: A Journal of Neurology 144 (6): 1853–68. 10.1093/brain/awab061.

Vanni, Silvia, Arianna Colini Baldeschi, Marco Zattoni, and Giuseppe Legname. 2020. ‘Brain Aging: A Ianus-Faced Player between Health and Neurodegeneration’. Journal of Neuroscience Research 98 (2): 299–311. 10.1002/jnr.24379.

Wang, Lei, Linda Yang, Marcella Debidda, David Witte, and Yi Zheng. 2007. ‘Cdc42 GTPase-Activating Protein Deficiency Promotes Genomic Instability and Premature Aging-like Phenotypes’. Proceedings of the National Academy of Sciences of the United States of America 104 (4): 1248–53. 10.1073/pnas.0609149104.

Wang, Qi-Min, Yu-Yu Xu, Shang Liu, and Ze-Gang Ma. 2017. ‘Isradipine Attenuates MPTP-Induced Dopamine Neuron Degeneration by Inhibiting up-Regulation of L-Type Calcium Channels and Iron Accumulation in the Substantia Nigra of Mice’. Oncotarget 8 (29): 47284–95. 10.18632/oncotarget.17618.

Ximerakis, Methodios, Scott L. Lipnick, Brendan T. Innes, Sean K. Simmons, Xian Adiconis, Danielle Dionne, Brittany A. Mayweather, et al. 2019. ‘Single-Cell Transcriptomic Profiling of the Aging Mouse Brain’. Nature Neuroscience 22 (10): 1696–1708. 10.1038/s41593-019-0491-3.

Young, Matthew D, and Sam Behjati. 2020. ‘SoupX Removes Ambient RNA Contamination from Droplet-Based Single-Cell RNA Sequencing Data’. GigaScience 9 (12): giaa151. 10.1093/gigascience/giaa151.

Zeisel, Amit, Hannah Hochgerner, Peter Lönnerberg, Anna Johnsson, Fatima Memic, Job van der Zwan, Martin Häring, et al. 2018. ‘Molecular Architecture of the Mouse Nervous System’. Cell 174 (4): 999–1014.e22. 10.1016/j.cell.2018.06.021.

Zheng, Grace X. Y., Jessica M. Terry, Phillip Belgrader, Paul Ryvkin, Zachary W. Bent, Ryan Wilson, Solongo B. Ziraldo, et al. 2017. ‘Massively Parallel Digital Transcriptional Profiling of Single Cells’. Nature Communications 8 (1): 14049. 10.1038/ncomms14049.

Zou, Jingjng, Kun Zhang, Jinde Zhu, Chaoyong Tu, and Jingqiang Guo. 2024. ‘Identification of Therapeutic Targets and Prognostic Biomarkers of the Ephrin Receptor Subfamily in Pancreatic Adenocarcinoma’. The Journal of International Medical Research 52 (1): 3000605231218559. 10.1177/03000605231218559.

