## Supplementary figures and images for "Molecular aging is the main driver of Parkinson’s Disease"

### Supplement Figure 1

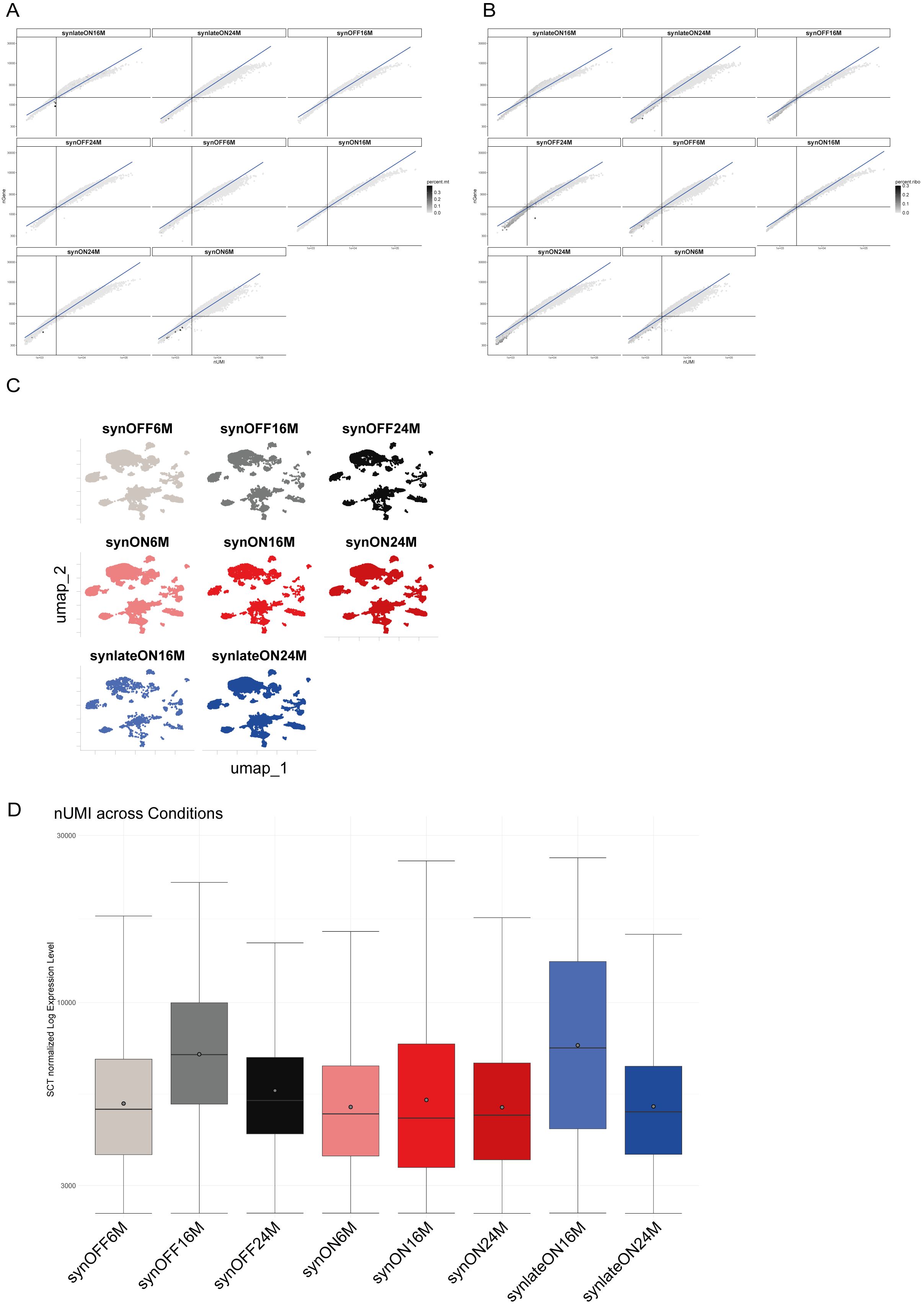

### Supplement Figure 2

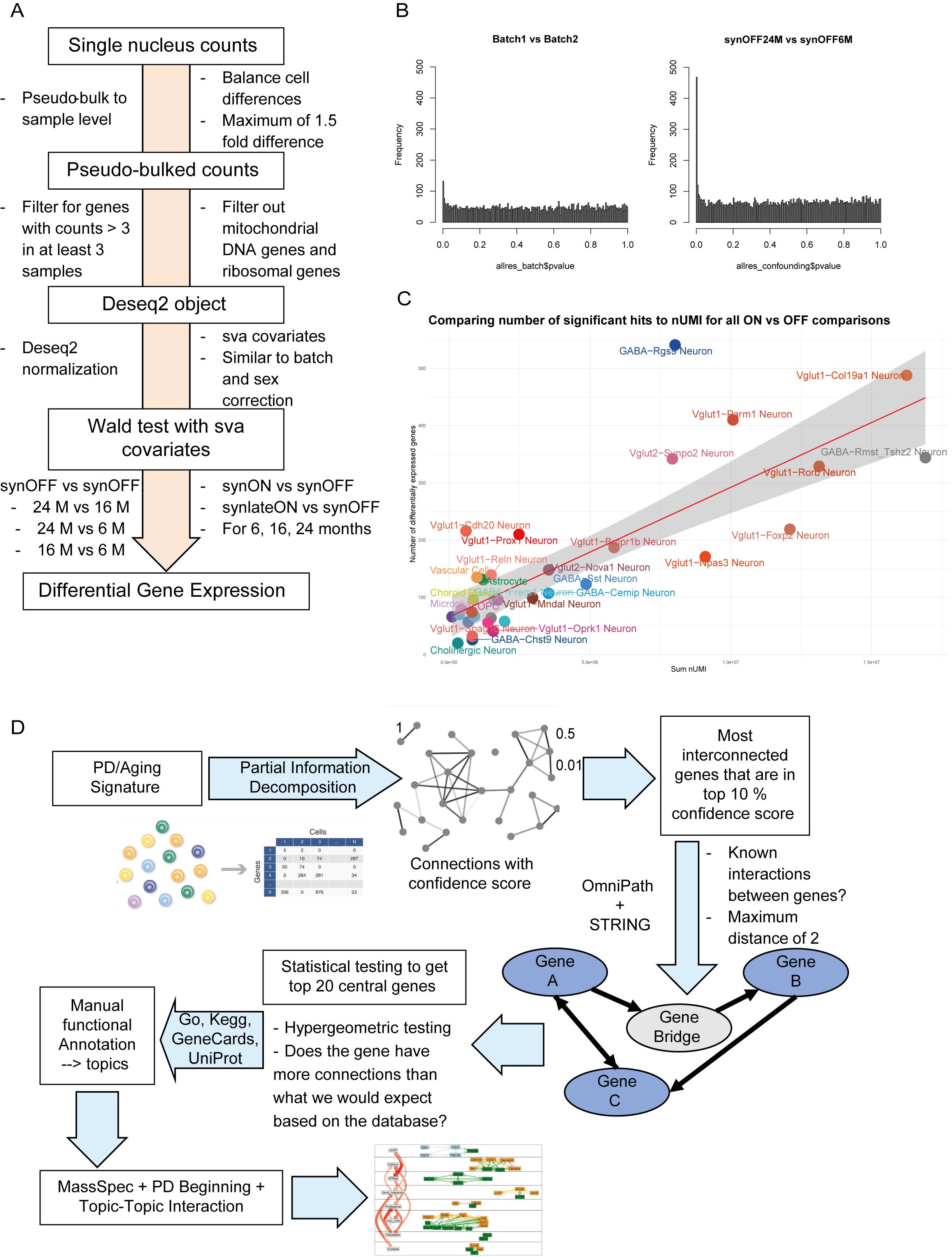

### Supplement Figure 3

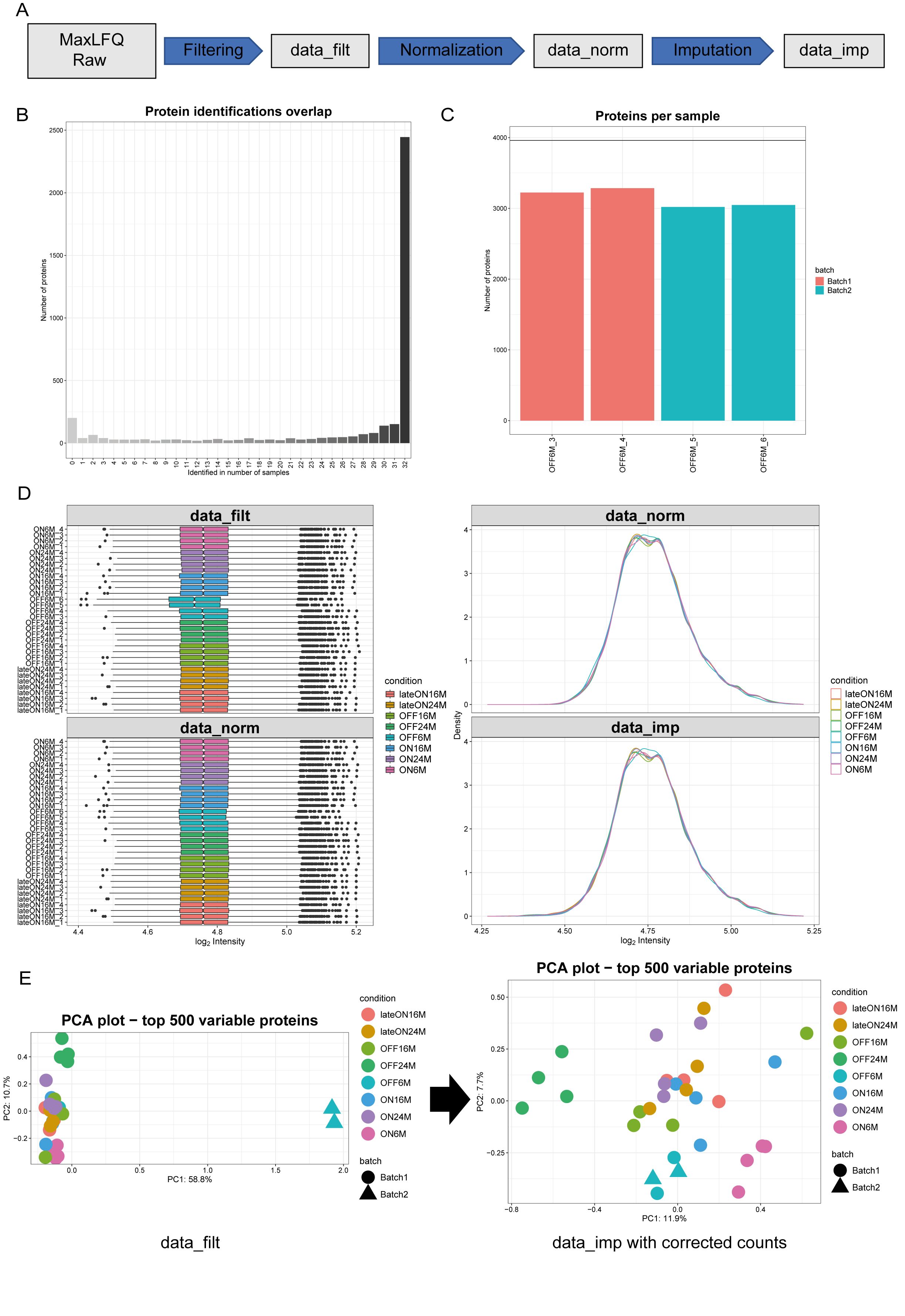

### Supplement Figure 4

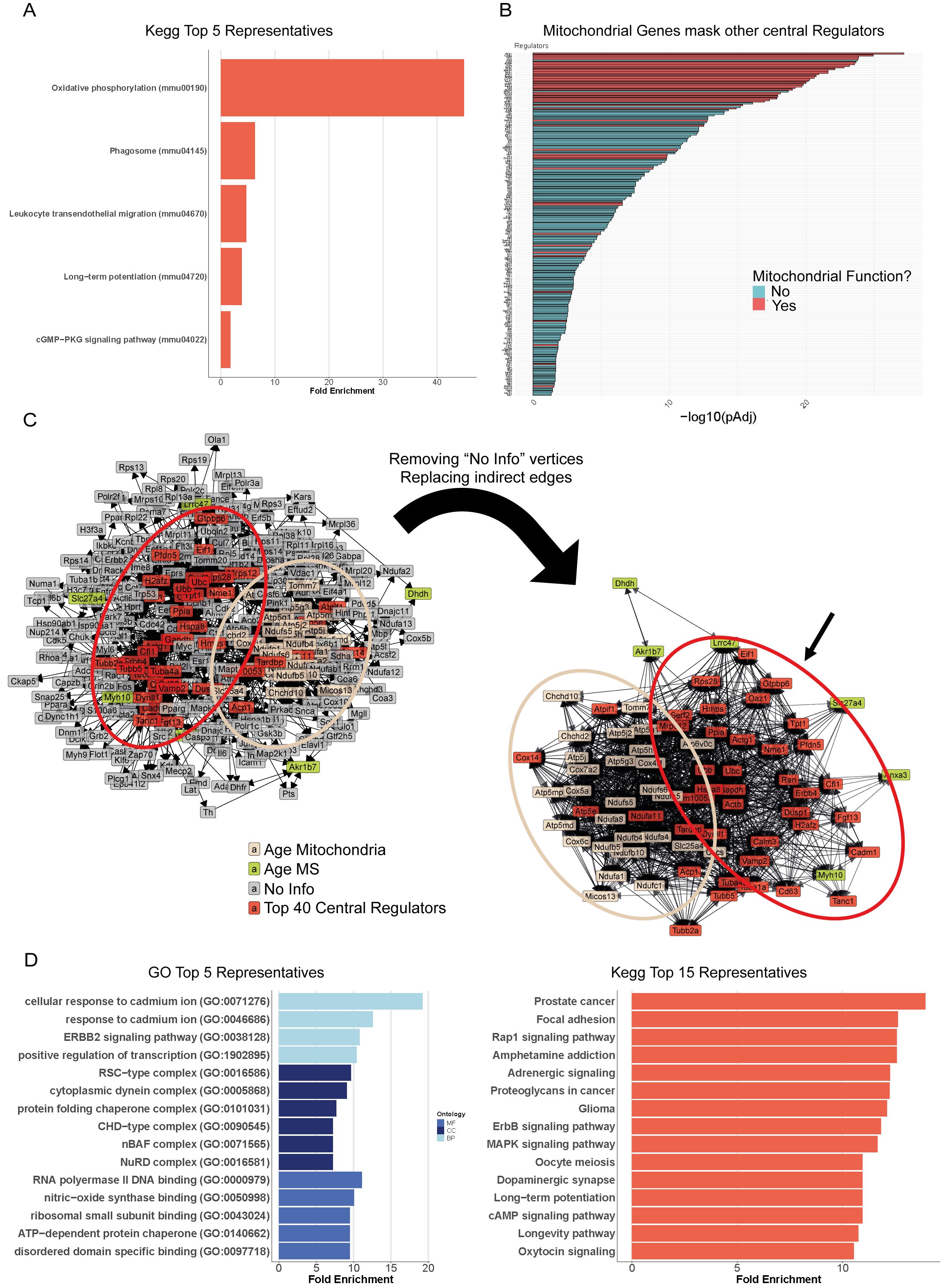

### Supplement Figure 5

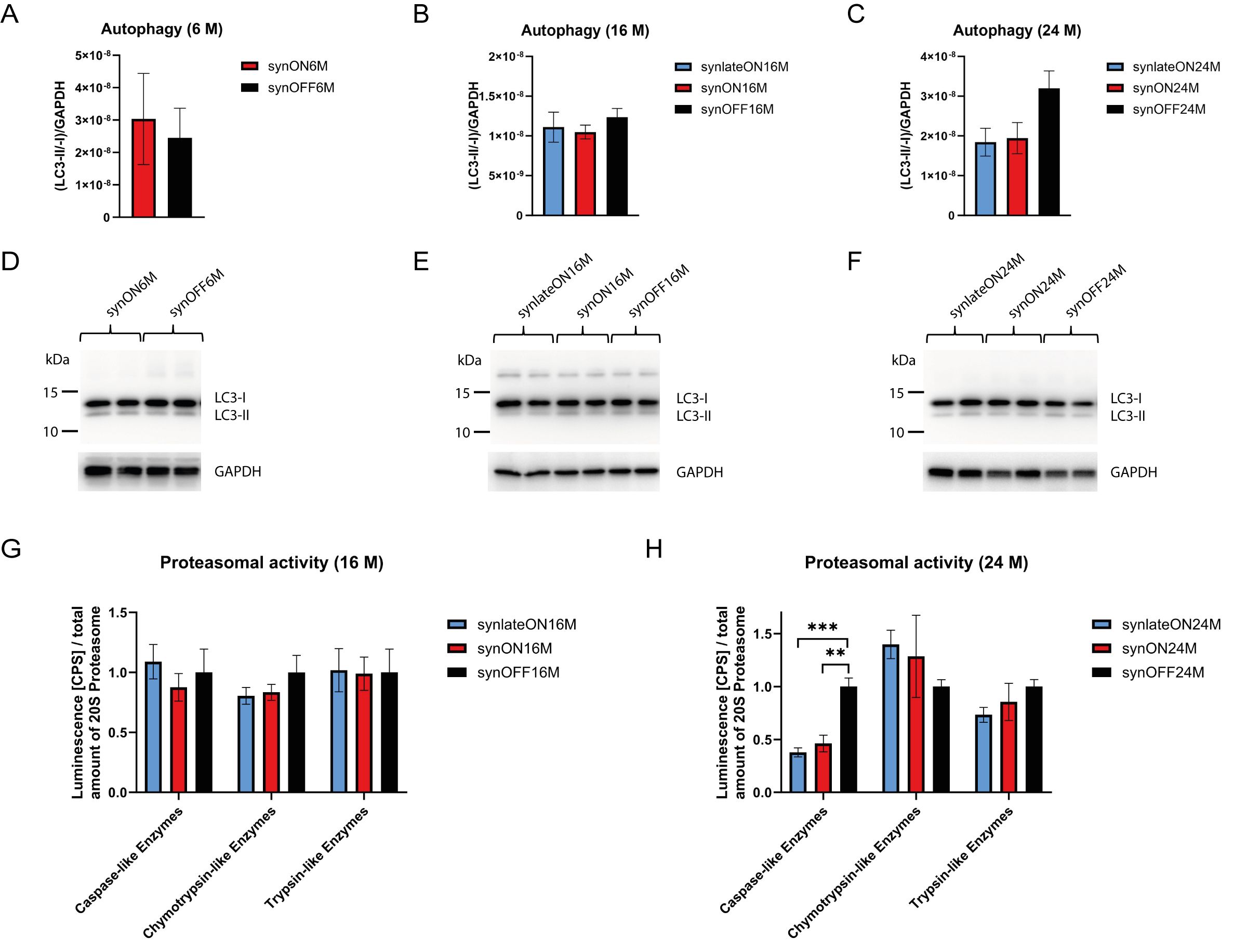

### Supplement Figure 6

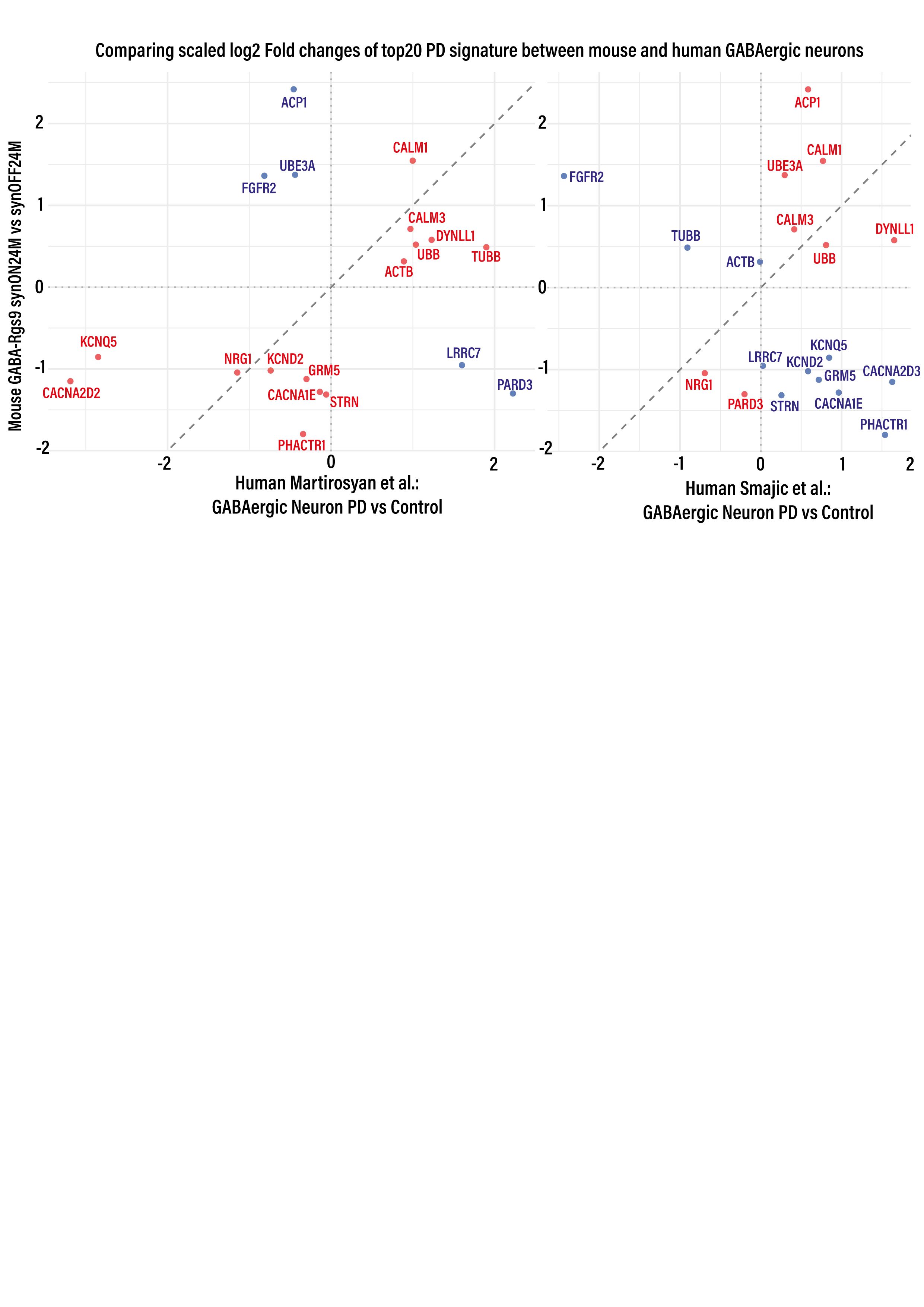
